# Cost-saving, trading, and internalization of externality: three economic strategies underlying amino acid archetypes of human metabolic enzymes

**DOI:** 10.64898/2026.03.25.714160

**Authors:** Kaixiang Yang, Jie Wei, Kai Sun, Hu Zhu, Yu Liu, Ziwei Dai

## Abstract

Amino acid composition of proteome reflects principles of metabolic resource allocation. While amino acids with higher biosynthetic costs are used less frequently in the proteome of unicellular organisms, whether this pattern applies to complex multicellular life remains elusive. Here we show that amino acid composition of human metabolic enzymes does not follow a simple cost-minimizing mode but instead segregate into four distinct amino-acid composition modules each associated with specific metabolic functions, a phenomenon we term emergence of amino acid archetypes (AArchetypes). The emergence of AArchetypes was associated with organismal complexity across a taxonomically diverse panel, indicating a shift in amino acid economics in complex organisms. These AArchetypes could be explained by three distinct economic strategies: cost-saving in young, fast-evolving enzymes; trading in conserved and widely expressed central carbon metabolic enzymes whose amino-acid compositions align with dietary intake; and internalization of negative externalities in intrinsically disordered proteins through elevated biosynthetic costs of IDR-related amino acids. In summary, this study develops a multi-layered economic framework that integrates production, exchange, and regulation to explain the diversification of amino acid composition in human metabolic enzymes.

## Introduction

Amino acids, fundamental building blocks of proteins, are not only symbols for encoding protein function but also physical entities that must be paid for with carbon, energy, and enzymatic capacity. *De novo* synthesis of amino acids requires three resources: carbon and nitrogen sources that provide the molecular backbone, energy that facilitates overcoming free energy barriers, and metabolic enzymes that catalyze reactions. The need to save costs of amino acid synthesis creates a simple economic law governing the organization of proteome in unicellular organisms: expensive amino acids are rare, and cheap ones dominate the proteome. The trade-off between cost-saving and maximizing protein sequence diversity produces a nearly universal negative linear relationship between costs of amino acids and logarithms of their frequencies in proteomes across bacteria and unicellular eukaryotes such as *E*.*coli, B. subtilis*, and *S*.*cerevisiae*^1-4^, as well as cancer cells^5^. This cost-frequency law has proven remarkably robust to environmental factors such as temperature^6^, redox potential^7^, and ocean depth^8^ that are known to affect amino acid composition of proteome.

Nevertheless, this law becomes conceptually fragile in complex organisms such as humans. Unlike bacteria, archaea and unicellular eukaryote, humans cannot synthesize all standard amino acids and must rely on dietary intake. Recent comparative analyses suggest that energetic selection contributed to the evolutionary outsourcing of costly amino acid biosynthesis in animals^9^. In human body, amino acids are obtained through diet, buffered by circulation, and redistributed across tissues with diverse metabolic and functional demands. Consequently, the cost of *de novo* synthesis no longer reflects the true economic price of an amino acid. Matching dietary amino acid compositions to cellular proteome has beneficial outcomes in multicellular animals such as fruit fly^10, 11^, implying an economic strategy of trading instead of production. Furthermore, the difference in amino acid synthesis pathways between multicellular and unicellular organisms^12^ could also profoundly change the costs of *de novo* amino acid synthesis. However, whether dietary supply differentially matches the compositional requirements of specific metabolic pathway groups remains unclear.

Furthermore, beyond the cost-frequency coupling of amino acids at the whole proteome level, a proteome is fundamentally not a single economic agent. Instead, it is a collection of proteins each serving distinct functions, operating in different cellular compartments, and facing different evolutionary and physiological constraints. Consequently, the cost-frequency law at proteome level does not constrain amino acid composition of single proteins. It operates at the protein abundance, where proteins with costly amino acid compositions have lower expression levels^13^. However, this observation does not address the important question of why such costly amino acid composition profiles still exist in a variety of proteins.

In this study, we focus on metabolic enzymes that represent a substantial fraction of cellular proteome to investigate principles governing the diversification of amino acid compositions of proteins. Here we show that, in organisms with greater complexity such as humans, the cost-frequency law does not simply weaken. Instead, it diverges into three distinct economic strategies. Four representative patterns of amino acid composition emerge in human metabolic enzymes, each enriched for specific biological functions and reflecting a specific strategy for managing metabolic cost. We refer to this phenomenon as emergence of amino acid archetypes (AArchetypes) and showed that AArchetypes tend to emerge in organisms with greater biological complexity.

By linking human AArchetypes to protein age, function, and structural properties, we show that these AArchetypes can be explained by three economic strategies: first, cost-saving, adopted by fast-evolving, young proteins to minimize synthetic costs of amino acids; second, trading, whereby conserved and widely expressed central carbon metabolic enzymes align their amino acid composition with dietary amino acid profiles; third, internalization of externality, in which intrinsic disordered region (IDR)-related amino acids are costlier, reflecting the possibility of “taxing” IDR-related amino acids to achieve effective regulation of IDR-related processes. These findings together establish an economic interpretation of proteomic amino acid composition in complex organisms by a layered economic system consisting of diverse economic strategies.

## Materials and methods

### Amino acid composition of metabolic enzymes and pathways

Information of metabolic pathways (i.e. KEGG pathways under the category “Metabolism”) was retrieved from the KEGG Pathway database. For a broad collection of species selected from KEGG Organisms to provide as comprehensive coverage as possible across domains, kingdoms, and phyla of Eukaryota, Bacteria, and Archaea, species-specific lists of genes and proteins encoding metabolic enzymes in each KEGG metabolic pathway and amino acid sequences of these enzymes were retrieved using the KEGG API. Relative frequency of each of the 20 standard amino acids across the full-length sequence (i.e., the residue count of a given amino acid divided by the total sequence length) was then calculated for each metabolic enzyme. The pathway-level amino acid composition was defined as the arithmetic mean of enzyme-level composition vectors across all enzymes annotated to the pathway.

### Calculation of amino acid enrichment scores for metabolic pathways

For each species, all metabolic enzymes were aggregated into a background metabolic enzyme pool. For each pathway *k* and amino acid *a*, we defined the observed pathway-level statistic *T*_obs_(*k,a*) as its pathway-level amino acid composition. We then performed a permutation test to quantify deviation from the background. Specifically, while keeping the number of enzymes in each pathway fixed, we repeatedly sampled the same number of enzymes from the background metabolic enzyme pool and computed the corresponding permuted statistic *T*_perm_(*k,a*). This procedure was repeated 1,000 times to simulate a null distribution. The empirical one-sided p value was then defined as follows:

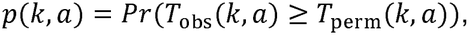

estimated by the fraction of permutations satisfying *T*_obs_(*k,a*) ≥ *T*_perm_(*k,a*) . Under this definition, large *p* values (*p* > 0.94) indicate enrichment (higher-than-expected amino acid fraction), whereas small p values (*p* < 0.05) indicate depletion (lower-than-expected fraction) relative to the background pool.

To represent enrichment/depletion patterns on a unified scale, we transformed p values into an AA enrichment score: p values within the non-significant interval (0.05 ≤ *p* ≤ 0.95) were set to 0.5 (neutral), whereas tail p values (*p* > 0.95 or *p* < 0.05) were kept unchanged. This yielded an AA enrichment score matrix (rows: pathways; columns: the 20 standard amino acids).

### Identification of AArchetypes

We performed hierarchical clustering of pathways based on Euclidean distance using the AA enrichment score matrix to identify candidate AArchetypes. For human, the metabolic pathways were clustered into four clusters. For other species, the number of clusters was determined by comparing distributions of silhouette coefficients across different numbers of clusters and selecting the number yielding highest average silhouette coefficient. KEGG functional category enrichment was performed for each module in each species using one-sided Fisher’s exact test to assess whether the module is enriched with major categories of metabolic functions such as carbohydrate metabolism, amino acid metabolism, and so on. Two metrics were considered in combination to identify AArchetypes in each species:

1. Proportion of significant AA enrichment/depletion. It is defined as the proportion of significant AA enrichment or depletion scores (i.e. p(k,a)<0.05 or p(k,a)>0.95) within the AA enrichment score matrix. This proportion assesses the extent of diversification in amino acid composition across different metabolic pathways.
2. Max(min enrichment p-value). For each pathway cluster, we calculated enrichment p-values for all KEGG functional categories and retained the minimum p-value. The maximum of these minimum p-values across all clusters was then used to assess whether every cluster was significantly enriched for at least one functional category.

Species with a max(min enrichment p-value) below 0.05, and a proportion of significant AA enrichment/depletion greater than 0.3 were considered to exhibit the emergence of AArchetypes.

### Random forest classification of human enzymes across AArchetypes

To test whether the four human AArchetypes exhibit separable amino acid composition profiles at the protein level, we trained a random forest classifier using the amino acid composition of human metabolic enzymes as features and AArchetype assignments as labels. Classification performance was evaluated in a one-vs-rest setting for each AArchetype, in which each AArchetype was considered in turn as the positive class, while proteins from the remaining three AArchetypes were pooled as the negative class. Discriminative ability was quantified by the area under the receiver operating characteristic curve (AUC).

### Correlation between amino acid composition and structure of human metabolic enzymes

To quantify the relationship between amino acid composition and 3D structure of human metabolic enzymes, we downloaded predicted structures (PDB format) from the AlphaFold database^14^. We then used Foldseek^15^ to convert each enzyme structure into a length-matched per-residue structural sequence, represented by a 20-state structural alphabet that encodes local structural environments along the polypeptide chain. Pairwise Euclidean distances between these structural sequences were computed using DECIPHER^16^, generating a structural distance matrix. We then computed pairwise Euclidean distances between amino acid composition vectors of the enzymes to obtain an amino acid composition distance matrix. Pearson correlation between these two distance matrices was then calculated over all off-diagonal elements.

### Acquisition of proteome, amino acid exchange, and evolutionary datasets

Subcellular localization annotations for human metabolic enzymes and amino acid sequences of all proteins in *H. sapiens* and *S. cerevisiae* were downloaded from UniProt. For each species, we computed the proteome-level abundance of the 20 standard amino acids, defined as the average amino acid composition across all proteins in the proteome. Human proteins were dichotomized into membrane-associated proteins and non-membrane proteins based on UniProt membrane-related annotations. Protein abundance data in human tissue samples, including breast, skin, liver, lung, and kidney, were obtained from ProteomicsDB^17^ (https://www.proteomicsdb.org/). Protein abundance datasets in human cell lines were obtained from CCLE^18^ (https://sites.broadinstitute.org/ccle/). We computed the average abundance of each protein across tissues to get its mean expression level. Protein abundance data for *S. cerevisiae* were obtained from a previous study by Brandon et al^19^. We used the mean protein abundance (molecules per cell) in the dataset as the expression level of yeast protein.

To quantify tissue specific expression, we calculated the Gini coefficient of tissue resolved protein abundance for each enzyme (higher Gini indicates stronger tissue specificity). Given the expression levels of a protein across *n* tissues *x*_1_,*x*_2_,…, *x*_*n*_ its mean expression level *µ* is:

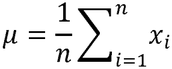

The Gini coefficient can be defined as the normalized mean absolute difference between all pairs of values:

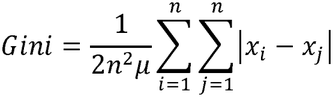

Differences in protein abundance and Gini coefficients between AArchetypes were evaluated using two-sided Wilcoxon rank-sum tests.

Metabolite uptake and secretion datasets in NCI-60 cell lines were obtained from a previous study by Aaron et al^20^ to quantify the exchange fluxes of amino acids in human cells. Gene age data were obtained from the GenTree database^21^, and evolutionary rate data were retrieved from the review by Zhang et al^22^.

### Comparison of human metabolic pathways with dietary patterns and food categories

Amino acid composition profiles for ten human dietary patterns and food-level amino acid contents with predefined category labels were obtained from our previous study^23^. For each food category, we calculated the arithmetic mean of amino acid measurement across its constituent foods and normalized the resulting category-level vector to sum to 100. Because experimental quantification of amino acids in foods often cannot reliably distinguish glutamate from glutamine, or aspartate from asparagine, fractions of these pairs of amino acids in human enzymes were merged before being compared. To match the combined measurements in dietary and food data, we merged the corresponding components in the pathway profiles. All comparisons used the resulting 18-component percentage vectors.

We performed t-distributed stochastic neighbor embedding (t-SNE; perplexity = 18) on the combined human pathway and dietary-pattern profiles. Euclidean distances were calculated for each pathway–dietary-pattern pair and summarized by the median. For each AArchetype A pathway, we separately calculated the median distance to dietary patterns and food categories. Their difference was defined as the food-category median minus the dietary-pattern median, and the paired medians were compared across pathways using a two-sided Wilcoxon signed-rank test.

### Cross-species comparisons of pathway and dietary amino acid composition

We matched pathways in human AArchetypes A–D to species-specific clusters in *Arabidopsis thaliana, Coffea arabica*, and *Solanum tuberosum* using KEGG pathway identifiers. Each distinct shared pathway–human AArchetype–plant cluster assignment was counted once, including pathways with multiple assignments. For dietary comparisons, we used Cluster 1 of each plant species with the same amino acid components and Euclidean distance calculation. For each group–dietary-pattern pair, distances were summarized by the median across pathways. Principal component analysis(PCA) of the combined pathway and dietary-pattern profiles was performed after centering and scaling each amino acid component to unit variance.

For the broader cross-species comparison, we identified pathways sharing a KEGG identifier with a human AArchetype A pathway. We calculated the median distance to the ten human dietary patterns for each pathway, then took the median of these pathway-level values for each species. Human reference profiles included AArchetype A-assigned pathways and trophic modes were assigned from taxonomic groupings with manual assignments for selected species. For species-specific comparisons, we used amino acid profiles for the *Drosophila melanogaster* standard diet (STD)^24^, the *Danio rerio* Unhydro, 50% Hydro, and 100% Hydro diets^25^, and the *Mus musculus* Chow, Casein, and Whey diets^26^. After aligning these profiles to the same 18-component percentage scale, we linked pathway compositions to species-specific cluster assignments by KEGG identifier and calculated each pathway–diet distance. A pathway was classified as human AArchetype A-matched when its KEGG identifier matched a human pathway containing at least one AArchetype A-assigned gene.

### Calculation of protein cost and energy cost for amino acids

We quantified the protein and energetic costs required for the biosynthesis of non-essential amino acids under a physiologically relevant, high-growth condition. Specifically, we considered a nutrient-replete environment in which water, oxygen, inorganic salts, glucose, amino acids, and other essential nutrients (Table S2) are available. First, we used GECKO^27^ to augment a human genome-scale metabolic model (Recon3D) with enzyme-usage constraints for individual reactions. For each metabolic enzyme, we computed the ratio of its molecular weight to its catalytic constant (*Mw*/*k*_*cat*_ ) to provide a proxy for the steady-state enzyme mass required to sustain one unit of metabolic flux through reactions catalyzed by it. For each metabolic reaction, a reaction-level protein cost coefficient *c*_*p,r*_ was computed by the minimal *Mw*/*k*_*cat*_ among all isozymes catalyzing this reaction. This reaction-level protein cost coefficient quantifies the minimal mass of enzyme required to carry one unit of flux through this reaction.

We also defined a reaction-level energy cost coefficient *c*_*e,r*_ of each reaction, quantified by the number of ATP molecules hydrolyzed in this reaction. Let *v*_*r*_ denote the flux through reaction *r*, the protein and energy costs consumed by this reaction can be calculated by *c*_*p,r*_ *v*_*r*_ and *c*_*e,r*_ *v*_*r*_, respectively. Collecting all reaction-level coefficients into vectors *c*_*p*_ and *c*_*e*_, the total protein cost and energy cost corresponding to a metabolic flux configuration *v* are 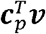 and 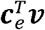, respectively.

We next applied flux balance analysis (FBA) to compute the marginal protein and energy costs associated with the biosynthesis of each non-essential amino acid. We formulated the problem as solving an optimal flux configuration *v* minimizing the total protein cost 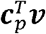 or energy cost 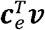 under flux balance, sustained growth, and other constraints. The following constraints were included in the optimization:

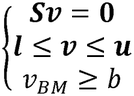

In which ***s*** is the stoichiometric matrix, ***l*** and ***u*** are lower and upper bounds of fluxes, *v*_*BM*_ is the biomass synthesis flux, *b* is a preset positive lower bound of the biomass synthesis flux. Default bounds of reactions were set to [0,1000] unless specified later. Exchange reactions (i.e. reactions with the form “⇒X[e]”, representing importing a metabolite from the external environment into the extracellular space) for molecules other than the growth-supporting nutrients listed in Table S2 were disabled by setting both their lower and upper bounds to 0. Upper bounds for exchange reactions for growth-supporting nutrients in Table S2 were set by referencing nutrient intake under human dietary patterns^28^.

We first computed the maximal feasible biomass synthesis flux *v*_*BM*,max_ under these constraints by solving the linear programming problem below:

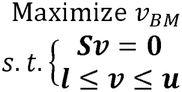

The lower bound of biomass synthesis flux was then set to 0.5*v*_*BM*,max_, thereby constraining growth to at least 50% of the maximal possible rate.

For each non-essential amino acid *A*_*i*_, upper bound of its uptake flux was set to 0 to enforce its *de novo* synthesis. We first computed the protein cost *P*_0_ and energy cost *E*_0_ at a reference state under the high-growth constraint:

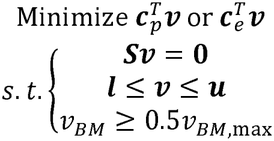

Next, to quantify the marginal cost of *A*_*i*_, a small positive number was added to its stoichiometric coefficient *α*_*i*_ in the biomass reaction to simulate an *A*_*i*_ - demanding state. We then recomputed the minimal protein and energy costs *p*_1(i)_ and*E*_1(i)_ at this *A*_*i*_ -demanding state by minimizing 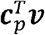 and 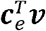. Finally, marginal protein and energy costs of the amino acid *A*_*i*_ were calculated as follows:

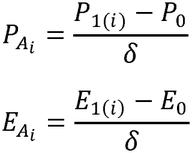

### Cost analysis of metabolic enzymes and pathways

For a metabolic enzyme with amino acid composition vector ***f***, in which *f*_*i*_ means the fraction of amino acid *A*_*i*_ in the sequence of that enzyme, its protein and energy costs were computed as follows:

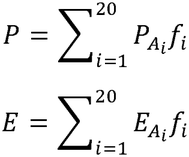

To evaluate how small changes in fraction of an individual amino acid affect the average protein cost of a metabolic pathway, we performed a cost sensitivity analysis for each pair of an amino acid and a metabolic pathway. Let ***f*** denote the average amino acid composition of enzymes in that pathway, ***P***_A_ and ***E***_*A*_ denote the vectors of protein and energy costs of amino acids, the protein and energy costs of this pathway were calculated by 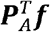 and 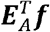, respectively. Next, to compute the protein and energy cost sensitivity score for amino acid *A*_*i*_, we randomly generated *n* amino acid compositions ***f***_1_’, ***f***_2_’, …, ***f***_*n*_’, where the fraction of *A*_*i*_ was set to ***f***_*n*_ + δ, δ is a small positive number. The cost sensitivity scores of *A*_*i*_ for this pathway (*S*_*P,i*_ for protein cost, *S*_*E,i*_ for energy cost) were then computed as follows.

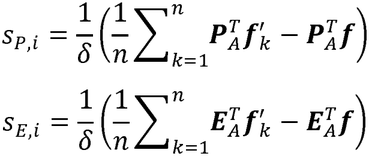

### IDR-associated analysis

Intrinsically disordered regions (IDRs) of proteins from human and other species were predicted using SPARROW^29^ . IDR proportion of a protein was calculated by dividing the total length of predicted IDRs within the protein by its total length. The four AArchetypes were further grouped into two pairs based on whether they are dominated by membrane proteins: AArchetypes A and B (non-membrane proteins), and AArchetypes C and D (membrane proteins). To test whether IDR-containing proteins (IDR proteins) are associated with higher protein cost at the proteome level, we carried out gene set enrichment analysis (GSEA) for the gene set. The human proteome was ranked by protein cost from high to low, and this ranked gene list was used as input for GSEA against KEGG pathways as well as the gene set corresponding to IDR proteins. The resulting normalized enrichment scores (NES) and visualization of their distributions were used to assess the enrichment.

### Acquisition of cellular fitness data upon gene overexpression

Cellular fitness data upon gene overexpression in human cells and *S. cerevisiae* were obtained from previous studies by Sack et al^30^ and Arita et al^31^, respectively. These datasets were used to investigate whether the overexpression of genes encoding proteins with higher IDR content is associated with reduced cellular fitness.

### Mathematical model for internalization of IDR externality

The model involves two groups of amino acids: an IDR-related amino acid group F with average cost of *X*, and an IDR-unrelated amino acid group *w* with average cost of 1, that together forms a protein with abundance *A*. Therefore, the total cost of the protein can be written as:

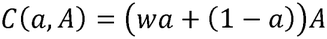

In which *a* is the fraction of the IDR-related amino acid group X in the protein. When the total biosynthetic cost of amino acids used in producing this protein is constrained by a budget (set as 1 for simplicity), we have an inequality constraint as follows:

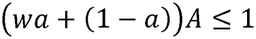

Next, we estimated the fraction of IDR in the sequence of this protein as a function *p*(*a*) by fitting a logistic function using sequences and IDR fraction of human proteins^32^ (see “Logistic regression model for IDR fraction” for details):

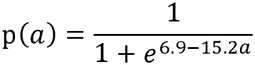

We therefore define a functional payoff function *P*_func_ (*a, A*) as the sum of the baseline benefit associated with the protein abundance A and the marginal increase of reward caused by IDR:

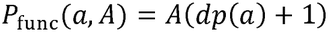

In which *d*, termed the IDR payoff gain, quantifies the increase in the payoff function contributed by one additional unit of IDR fraction. This functional payoff function quantifies how much the function of the protein is enhanced by IDRs in its sequence. The overall payoff function of the protein is then defined as the weighted sum of *P*_func_ (*a, A*) and an information capacity term *P*_info_(*a*) that quantifies the ability of the protein sequence to store information:

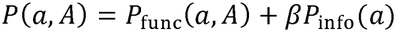

We assume that the protein determines its amino acid composition by maximizing the payoff function under the constraint of limited budget:

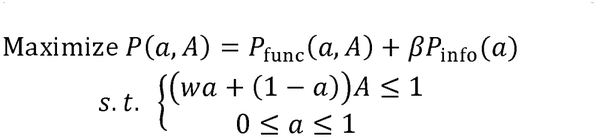

As *P*_func_ (*a, A*) is a linear function of the protein abundance *A* with the positive linear factor *dp*(*a*)+1, the payoff function *P*(*a, A*) is also a monotonically increasing function of *A*. Therefore, at the optimal solution, *A* equals to its maximally allowed value under the limited budget constraint:

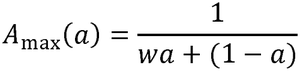

Therefore, the optimal payoff function can be computed by optimizing *a*:

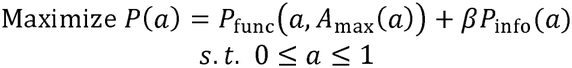

The constrained optimization problem was solved by the MATLAB built-in function “fmincon”.

### Logistic regression model for IDR fraction

We partitioned the 20 amino acids into two sets according to their association with IDR: X (IDR-related amino acids: M, K, R, S, Q, P, E) and Y (all remaining amino acids). For the *i*-th protein with amino acid composition *f*_*i*_, its fraction of IDR-related amino acids was calculated as follows:

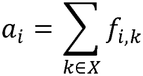

To model how the fraction of IDR depends on *a*, we fitted a binary logistic regression model with *a* as the predictor and the IDR-containing label *y* as the outcome:

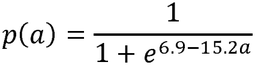

### Definition of information capacity term

The information capacity term of a protein was defined as a function *P*_info_(*a*) of fraction of IDR-related amino acids in the sequence of this protein, *a*. To derive the analytical expression of *P*_info_(*a*), we started from computing the Shannon entropy of a protein from its composition of 20 standard amino acids, ***f*** = (*f*_1_, *f*_2_, …, ***f***_20_):

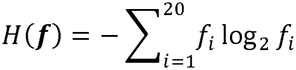

For a fixed sequence length, the logarithm of the number of sequences compatible with a given composition is proportional to *H*(***f***). The information capacity *P*_info_(*a*) was then defined as maximal *H*(***f***) among all possible sequences satisfying ∑_*i* ∈*X*_ *f*_*i*_ = a (i.e. total fraction of amino acids in the IDR-related group *X* equals to *a*):

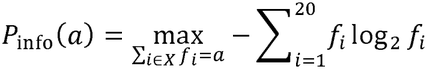

As group *X* contains 7 amino acids, the sequence achieves maximal Shannon entropy when the fraction *a* is uniformly distributed across the 7 amino acids in *X* (IDR-related amino acids) and the remaining fraction 1 − *a* is uniformly distributed across the 13 amino acids in *Y* (IDR-unrelated amino acids). Therefore, the information capacity corresponding to can be calculated:

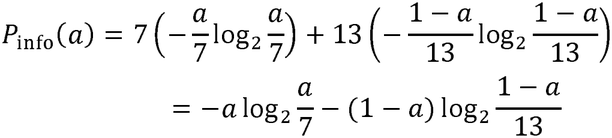

This function therefore quantifies how the size of the accessible sequence space varies as the fraction of IDR-related residues changes.

### Estimation of species-specific IDR cost ratio

For *H*. sapiens and *S*. cerevisiae, we first computed their average amino acid composition over the entire proteome. Let *X* and *Y* denote the groups of IDRrelated and IDR-unrelated amino acids, the IDR cost ratio G was defined as follows:

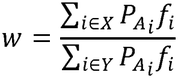

In which 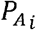 is the marginal protein cost of amino acid *A*_*i*_ in this species (human or yeast), *f*_*i*_ is the average fraction of amino acid *A*_*i*_ in proteins from this species.

### Inference of parameters *β* and *d*

The model parameter *β* was inferred based on the assumption that in yeast (*w* = 0.89) where proteins contain only a small fraction of IDRs, a protein with median *a* (*a* = 0.37) does not benefit from IDR. This assumption means that under the parameter set { *w* = 0.89, *d* =0}, the optimal solution of *a* is 0.37.

Hence, we sought to determine the parameter *β* that gives this optimal solution. We computed optimal values of *a* (referred to as *a*_opt_(*β*)) for an array of 1001 evenly distributed *β* values in the range of [0,10], {*β*_*i*_ = 0.01(*i* − 1)}, and then selected the two values *β*_*i*_ and *β*_*i* +1_ that satisfies *a*_opt_(*β*_*i*_) > 0.37 and *a*_opt_(*β*_*i* +1_) < 0.37. Assuming that *a*_opt_ changes linearly with *β* in the range [*β*_*i*_, *β*_*i* +1_], the parameter *β*corresponding to *a*_opt_ = 0.37 was computed as follows by inverse linear interpolation:

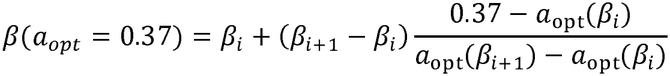

Similarly, to infer the parameter for a human protein with IDR-related amino acids fraction *a*, optimal values of *a* (referred to as *a*_opt_(*d*)) were computed for 3001 evenly distributed values of in the range of [0,30]. For each given *a* in the range [*a*_opt_(0),*a*_opt_(30)], its corresponding value of *d* was also computed using inverse linear interpolation.

## Results

### Emergence of AArchetypes in human metabolic enzymes

To test whether the amino acid composition of human proteins still follows a single cost-minimization principle, we focused on metabolic enzymes that constitute a large fraction of the human proteome^33^. We obtained amino acid sequences of all human metabolic enzymes from the KEGG database and quantified pathway-level amino acid composition by computing an amino acid (AA) enrichment score using permutation test (Fig. 1a, Methods). Hierarchical clustering of the amino acid enrichment scores of KEGG metabolic pathways identified four discrete clusters each characterized by a distinct pattern of amino acid usage (Fig. 1b). Significant enrichment of major metabolic categories was observed in each cluster: cluster A is enriched with carbohydrate metabolism, cluster B is enriched with amino acid metabolism, cluster C is enriched with xenobiotics metabolism, and cluster D is enriched with lipid and glycan metabolism (Fig. 1c). Furthermore, although the analysis was performed at the level of metabolic pathways, individual metabolic enzymes in each cluster can be distinguished from those in remaining clusters solely based on their amino acid composition (Fig. 1d), confirming that the diversification of amino acid composition occurs at the level of single proteins rather than being an artifact of pathway-level averaging.

**Fig. 1.**
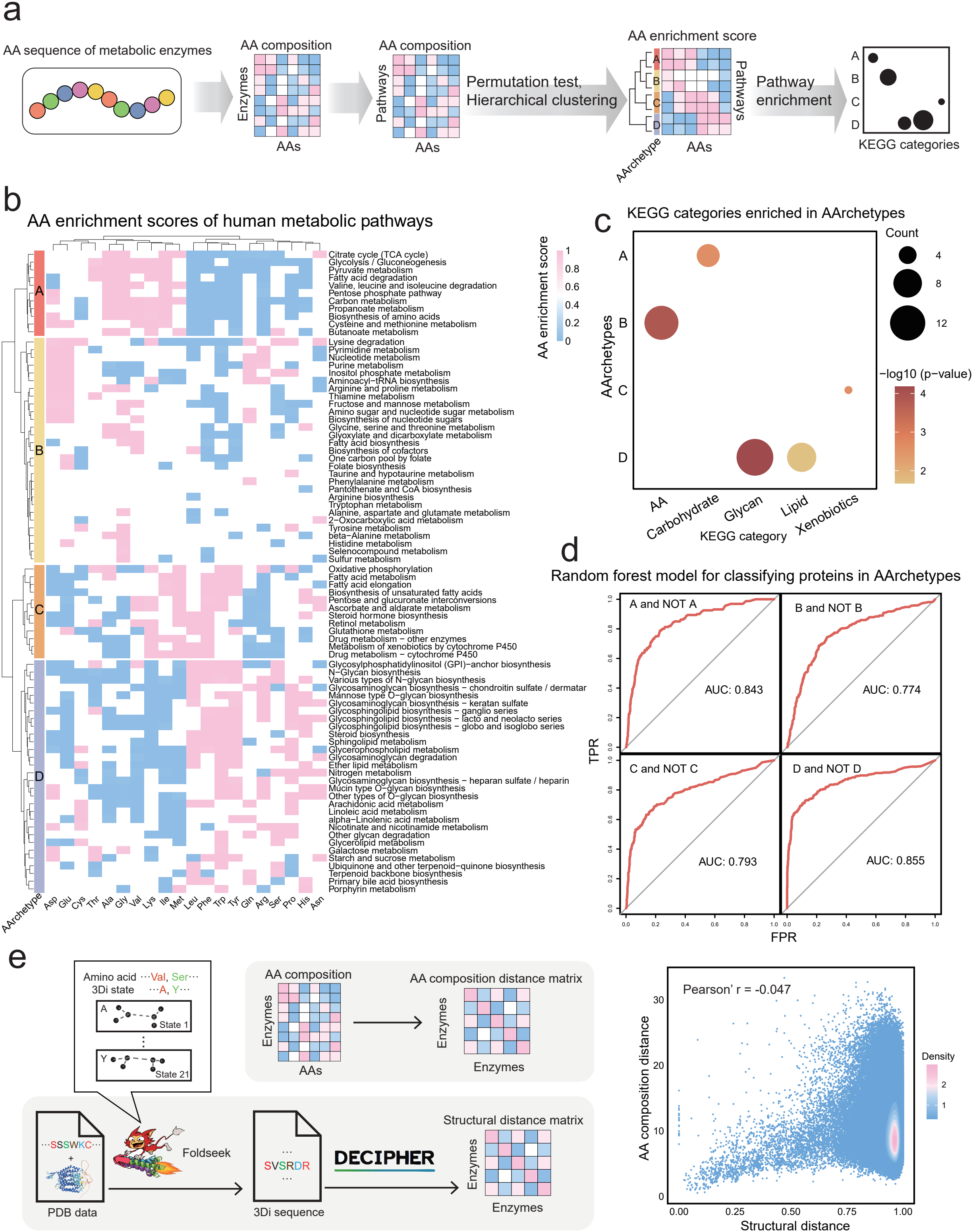
Emergence of AArchetypes in human metabolic enzymes. a: Workflow for calculating amino acid enrichment scores of metabolic pathways. b: Amino acid enrichment scores of human metabolic pathways and AArchetypes identified by clustering analysis. c: KEGG metabolic categories enriched in AArchetypes. d: Receiver operating characteristic (ROC) curves of random forest classifier to assess the separability of the four human AArchetypes using protein-level amino acid composition. e: Left: workflow for computing pairwise amino acid composition distance and structural distance matrices between proteins; Right: density scatter plot comparing amino acid composition distance to structural distance. Each dot represents a protein; colors indicate point density.

To test whether these clusters simply reflect structural similarity between functionally related proteins, we computed pairwise distances in amino acid composition with structural distances between metabolic enzymes (Fig. 1e, Methods). These two distances were poorly correlated (Pearson correlation = -0.047, Fig. 1e), suggesting that structural similarity does not explain the observed clustering of amino acid composition.

Together, these results reveal that human metabolic enzymes exhibit the emergence of four distinct amino acid archetypes (AArchetypes) each associated with specific metabolic functions. This observation indicates existence of factors beyond catalytic requirements or structural constraints that shape amino acid composition of human metabolic enzymes.

### AArchetype emergence is associated with greater organismal complexity

Next, we sought to investigate how the emergence of AArchetypes varies across the tree of life. To understand the emergence of AArchetypes from an evolutionary perspective, we selected 49 species from KEGG Organisms to achieve broad phylogenetic coverage across the tree of life and avoid dense sampling of a few closely related lineages (Fig. 2a, Methods). These selected species span all three domains of life and cover multiple major clades within each domain, encompassing both unicellular and multicellular organisms and phylogenetically diverse plant, fungal, and animal lineages. For each species, we computed pathway-level average amino acid compositions and performed classical multidimensional scaling (MDS) to visualize the global landscape of amino acid composition across species (Fig. 2b). Pathways from multicellular organisms tended to occupy a more compact region of the MDS space, whereas those from unicellular organisms were more broadly distributed (Fig. 2b). This difference motivated us to hypothesize that the observed emergence of AArchetypes in human metabolic enzymes could be associated with multicellularity and organismal complexity.

**Fig. 2.**
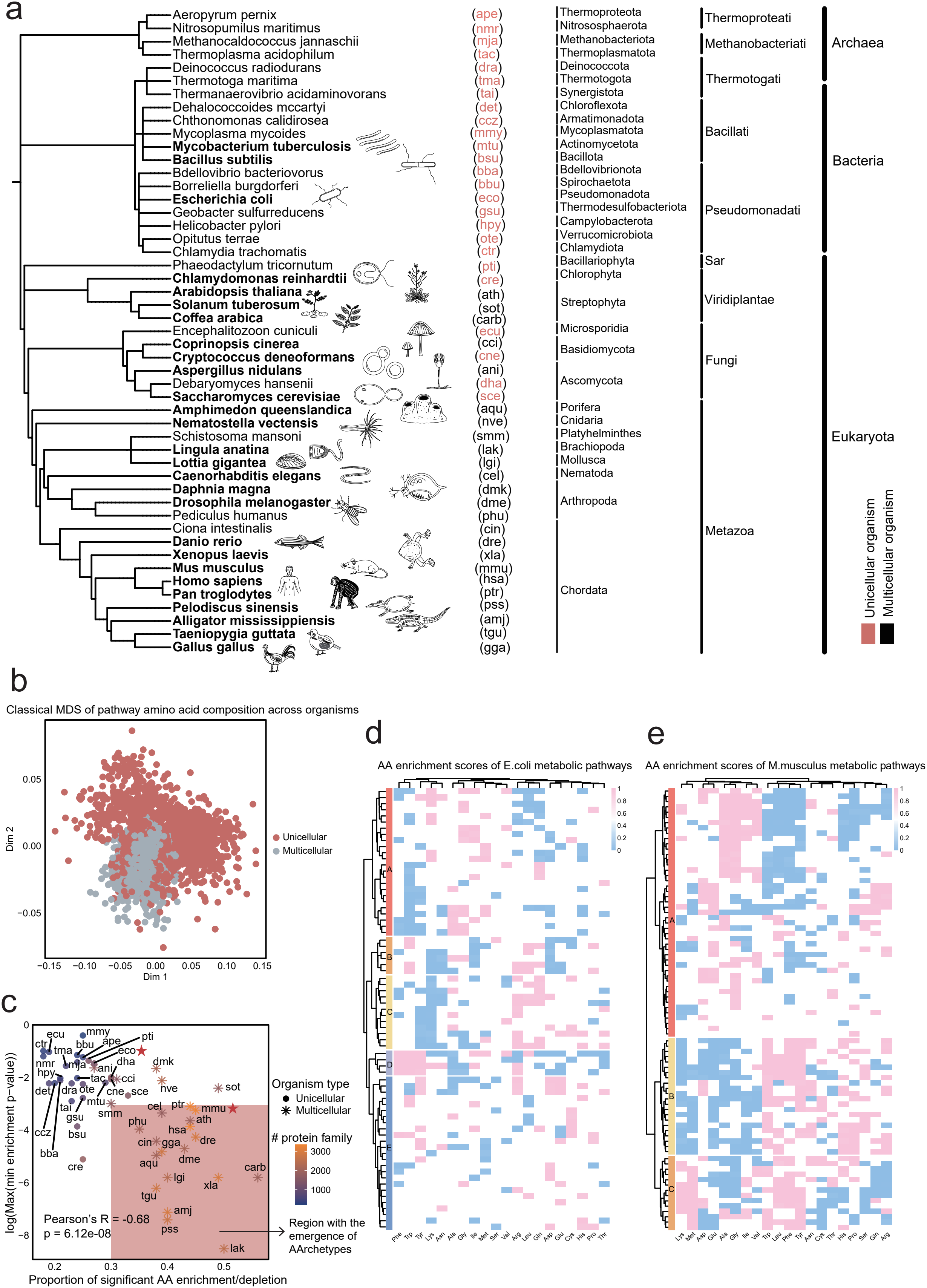
AArchetypes only emerge in multicellular organisms. a: Phylogenetic tree of 25 unicellular organisms and 24 multicellular organisms. In the phylogenetic tree, the labels from left to right indicate the full species name, the organism code in KEGG Organisms, and the phylum, kingdom, and domain in the taxonomic lineage. b: Classical multidimensional scaling (MDS) of pathway level amino acid composition arcoss all organisms. Each dot represents one metabolic pathway from one organism. c: Three independent metrics assesing the emergence of AArchetype in organisms. The AA enrichment scores for the two species marked with red stars (*eco* and *mmu*) are shown in Fig. 2d and e. A two-sided Pearson’s correlation test was used to compute the p-value. d, e: AA enrichment scores of metabolic pathways for *E*.*coli* and *M*.*musculus* (*eco* and *mmu*).

To test this hypothesis quantitatively, we applied the same enrichment and clustering framework used for human enzymes to all species (Fig. 2c–e and Fig. S1 and S2). We used two metrics to assess the presence of AArchetypes:

(1) proportion of significant AA enrichment or depletion, which quantifies the heterogeneity of amino acid usage across metabolic pathways; and (2) the maximal minimum (max–min) enrichment *P* value, which evaluates whether all clusters are enriched with metabolic categories. Together, these metrics assess whether metabolic enzymes in different pathways exhibit heterogeneous amino acid usage and whether the resulting clusters are functionally coherent.

Under this framework, we found that these two AArchetype-related features were significantly correlated across species: a higher proportion of significant AA enrichment or depletion is associated with a lower max–min enrichment *P* value (Pearson’s *R* = ™0.68, *P* = 6.12 × 10^™8^; Fig. 2c). Thus, stronger diversification of amino acid usage across metabolic pathways was accompanied by stronger functional enrichment of the resulting clusters.

Furthermore, emergence of AArchetypes was absent in a small subset of multicellular eukaryotes (such as *Aspergillus nidulans*, a filamentous fungus with relatively simple multicellular structure) and all unicellular species (Fig. S1), including unicellular eukaryotes, bacteria (Fig. 2d), and archaea. By contrast, nearly all multicellular eukaryotes examined exhibited clear emergence of AArchetypes (Fig. S2), including human (*Homo sapiens*, hsa), chimpanzee (*Pan troglodytes*, ptr), mouse (*Mus musculus*, mmu, Fig. 2e), fruit fly (*Drosophila melanogaster*, dme), worm (*Caenorhabditis elegans*, cel), and plant (*Arabidopsis thaliana*, ath). This observation implies an association between emergence of AArchetypes and multicellularity. We next asked whether the strength of AArchetype emergence also varies continuously against organismal complexity. To answer this question, we used the number of annotated protein families in each species as a proteome-level proxy for organismal complexity^34^. Species with greater number of protein families tended to show stronger signals of AArchetypes for both metrics, suggesting that AArchetypes preferentially emerge in organisms with greater complexity (Fig. 2c, S1, S2, Table S1). We next sought to understand what strategies and constraints could contribute to the divergence of amino acid composition across those AArchetypes.

### The youngest AArchetype exhibits the lowest amino acid synthetic cost

Given the well-established cost-minimizing principle in the proteomes of bacteria and unicellular eukaryotes, we asked whether biosynthetic cost might also contribute to the organization of amino acid usage across human AArchetypes. To characterize how amino acid economics differs across human AArchetypes, we computed the synthetic costs of all non-essential amino acids within the human metabolic network. Two types of costs were considered: protein cost, which quantifies the amount of enzymes needed for *de novo* synthesis of a unit of the amino acid, and energy cost, which quantifies the number of ATP molecules consumed in synthesizing a unit of the amino acid. Both costs were defined as marginal increases in total enzyme usage or ATP consumption upon a small increase in demand for each amino acid (Fig. 3a, S3a, Methods).

**Fig. 3.**
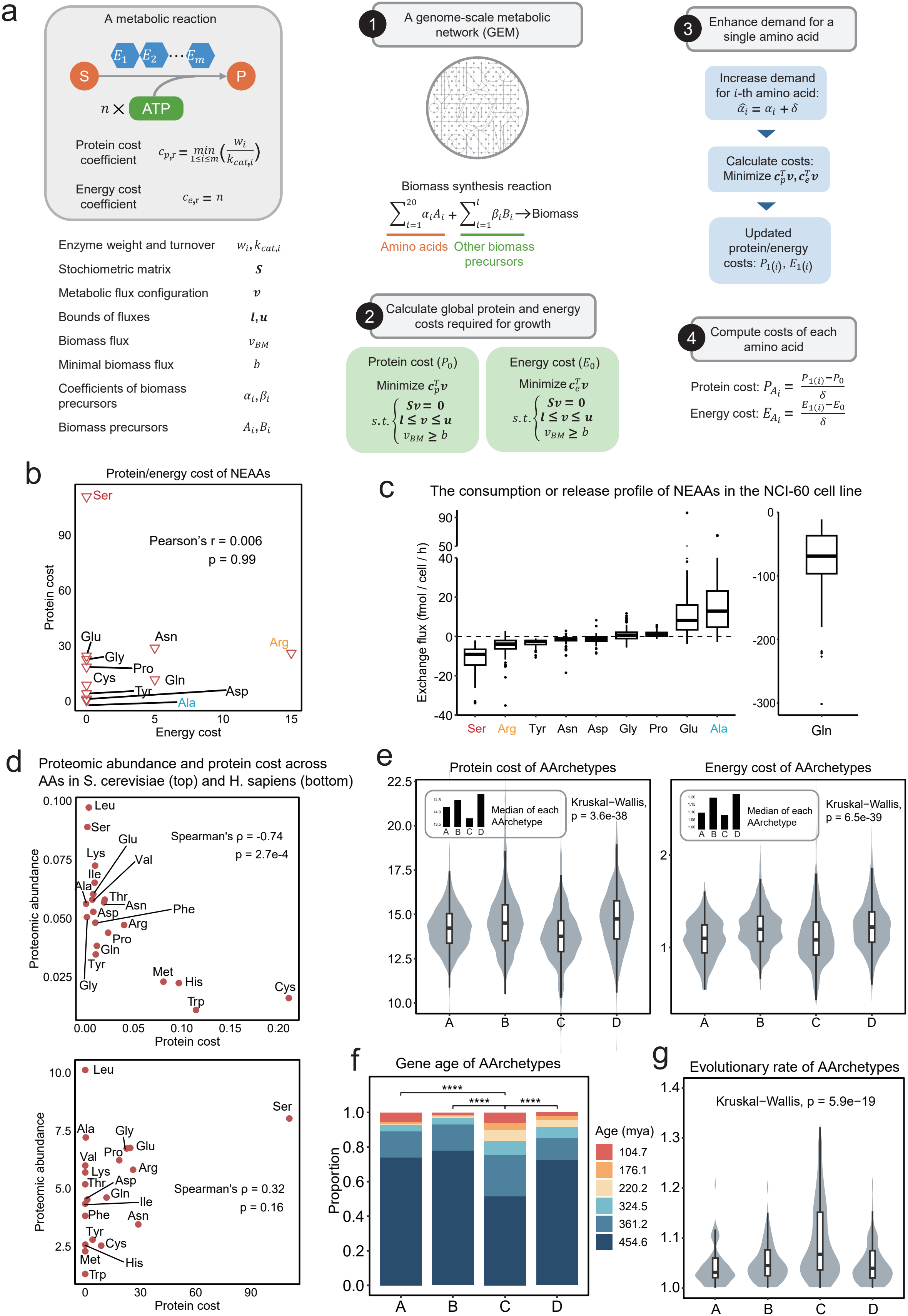
Youngest AArchetype minimizes synthetic cost. a: Schematic overview of the algorithm calculating protein and energy cost for amino acids. b: Protein cost and energy cost for non-essential amino acids in humans. A two-sided Pearson’s correlation test was used to compute the p-value. c: The consumption or release profile of non-essential amino acids in the NCI-60 human cell line panel. Among these, serine has the highest protein cost, arginine has the highest energy cost, and alanine has the lowest protein cost. d: Scatter plots comparing proteomic abundance and protein cost across amino acids in yeast (S. cerevisiae, top) and human (H. sapiens, bottom). A two-sided Spearman’s rank correlation test was performed to compute the p-values. e: Violin plots comparing protein cost (left) and energy cost (right) across the four human AArchetypes. P-values were computed using the Kruskal-Wallis test. Bar plots inside the panels show median values for the AArchetypes. f: Distribution of protein age among the four human AArchetypes. Statistical significance was evaluated using a Chi-square test of independence. g: Distribution of evolutionary rate for proteins in the four human AArchetypes. Statistical significance was evaluated using the Kruskal-Wallis test.

The two costs are uncorrelated at single amino acid level (Pearson correlation = 0.006, Fig. 3b), indicating that amino acids expensive in terms of enzymatic investment are not necessarily energy intensive, and *vice versa*. To examine whether these theoretical costs are consistent with amino acid utilization in human cells, we examined the uptake and secretion profiles of amino acids in the NCI-60 human cell lines. Amino acids with highest protein and energy costs, serine (highest protein cost) and arginine (highest energy cost), are also among those most actively taken up from the environment (Fig. 3c). This observation is consistent with the possibility that environmental uptake can reduce the need for *de novo* synthesis of costly amino acids. Applying the same framework to the budding yeast *S. cerevisiae* reproduced the known negative correlation^4^ between protein cost and proteome abundance of amino acids (Fig. 3d, top), therefore providing a consistency check for our cost estimates. However, although increase in synthetic cost is weakly correlated to decrease in proteome abundance in human compared to yeast (Fig. S3b), protein cost and proteome abundance of amino acids correlated poorly in human (Fig. 3d, bottom), suggesting that cost minimization alone could not fully explain the mode of amino acid usage in the human proteome.

We next asked whether cost minimization might operate selectively within specific AArchetypes. To our surprise, although protein and energy costs correlated poorly at single amino acid level (Fig. 3b), the four AArchetypes exhibit consistent ordering when these two costs were aggregated at the AArchetype level: AArchetype C is the most cost-effective, followed by AArchetype A, while AArchetypes B and D are the costliest (Fig. 3e). Thus, the four distinct AArchetypes exhibit systematic differences in amino acid synthetic cost that are consistent between two largely independent cost measures.

We next asked whether these cost differences were associated with the evolutionary history of enzymes. We estimated the age and evolutionary rate of human metabolic enzymes (Methods) and found that the most cost-effective AArchetype C was enriched with younger proteins (Fig. 3f) that evolve faster (Fig. 3g). Together, these results reveal an association between low amino acid synthetic cost and recent evolutionary origin. One possible explanation is that younger, faster-evolving enzymes have greater flexibility to adapt their amino acid compositions to achieve lower synthetic costs, while ancient and highly conserved enzymes that originated before the appearance of complex organisms might face stronger evolutionary constraints.

### Central carbon metabolism AArchetype matches dietary amino acid composition

Dietary intake is an important source of amino acids in animals including humans. Amino acids obtained from diet can reduce reliance on de novo synthesis and the associated biosynthetic costs^9^. Beyond cost-saving through biosynthetic optimization, enzymes whose amino acid composition is compatible with dietary supply may therefore obtain an effectively “cheaper” supply of amino acids, a strategy analogous to trading in economics. To examine this hypothesis, we compared the average amino acid compositions of metabolic pathways in each AArchetype with those of ten representative human dietary patterns^23^. Joint t-distributed stochastic neighbor embedding (t-SNE) projection indicated that dietary profiles appeared closest to AArchetype A pathways, which are enriched in carbohydrate and central carbon metabolism(Fig. 4a). Consistent with this visual pattern, AArchetype A exhibited the smallest median Euclidean distance to all dietary patterns among the four AArchetypes(Fig. 4b, Methods). The consistency across diverse dietary patterns suggests a broad compositional compatibility between dietary amino acid supply and central carbon metabolism enzymes. Such matching may be biologically relevant, as previous studies in model organisms such as fruit fly have reported beneficial health outcomes from proteome-matching dietary amino acid compositions^35^. To further evaluate the proposed trading strategy in AArchetype A, we next asked whether this compatibility was more pronounced for dietary patterns, which combine amino acids from multiple foods, than for individual food categories. We therefore compared their distances to dietary-pattern and food-category profiles.

**Fig. 4.**
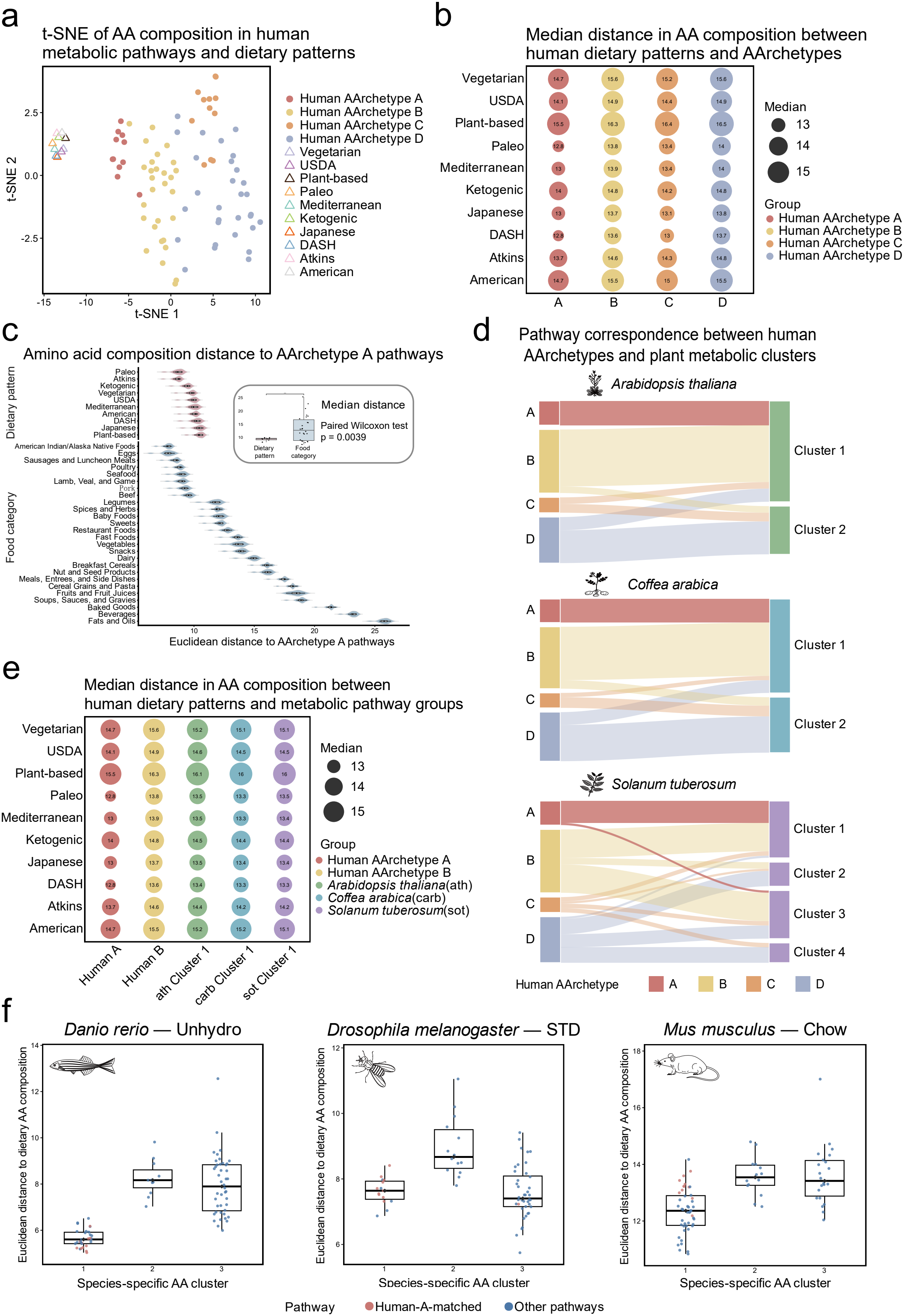
Dietary amino acid composition matching of central carbon metabolism AArchetype A across species. a: t-distributed stochastic neighbor embedding (t-SNE) of amino acid composition profiles of human metabolic pathways (circles for AArchetype A-D and triangles for ten human dietary patterns). b: Bubble plot showing the median Euclidean distance between amino acid compositions of each human dietary pattern and AArchetype A–D. c: Distributions of Euclidean distances between amino acid composition of individual AArchetype A pathways and human dietary patterns or food categories. The inset compares pathway-level median distances between dietary patterns and food categories using a paired Wilcoxon test (P = 0.0039). d: Sankey diagrams showing the correspondence of metabolic pathway assignments between human AArchetype A–D and species-specific metabolic pathway clusters in three plant species (*Arabidopsis thaliana, Coffea arabica*, and *Solanum tuberosum*). The width of each Sankey flow represents the number of shared pathway assignments connecting a human AArchetype to a plant cluster. e: Bubble plot showing the median Euclidean distance between amino acid compositions of human dietary patterns and human AArchetypes A and B or Cluster 1 metabolic pathways from *A. thaliana, C. arabica*, and *S. tuberosum*. f: Distributions of Euclidean distances between species-specific metabolic pathway clusters and corresponding experimental dietary amino acid compositions in *Danio rerio* (Unhydro), *Drosophila melanogaster* (STD), and *Mus musculus* (Chow).

AArchetype A pathways had smaller median distances to dietary patterns than to food categories (Fig. 4c, Methods), and this relationship held for every pathway in the group (Fig. S4a). These results support the possibility that combinations of foods in dietary patterns provide a closer amino acid composition to central carbon metabolism pathways than individual food categories do. Consistent with the potential benefits of such matching, exome-informed protein blends have been shown to improve growth and feed conversion efficiency in protein-limited mice^36^.

Broad similarities in exome-derived amino acid profiles have been reported across diverse taxa^37^, raising the possibility that dietary matching observed in humans partly reflects general compositional conservation. Autotrophic plants share conserved central carbon metabolic pathways with human but do not depend on dietary protein intake, providing a comparison for evaluating the proposed trading strategy. To establish a basis for comparing the corresponding metabolic groups, we compared human AArchetype assignments with plant species-specific clusters to first determine whether pathways corresponding to human AArchetype A were similarly concentrated within a single cluster in each plant species. Pathways assigned to human AArchetype A were predominantly found in Cluster 1 of *Arabidopsis thaliana, Coffea arabica* and *Solanum tuberosum* (Fig. 4d). The complementary PCA showed partial overlap in amino acid composition between human AArchetype A and these plant clusters, consistent with their shared pathway composition(Fig. S4b). We then compared human AArchetypes A and B and Cluster 1 of each plant with the ten human dietary patterns. Across all dietary patterns, Cluster 1 of each plant had greater median Euclidean distances than human AArchetype A(Fig. 4e). Thus, despite their correspondence in pathway membership, human AArchetype A consistently matched human dietary amino acid composition more closely than the plant clusters. A broader comparison of AArchetype A-matched pathways likewise showed that representative autotrophic species generally had larger distances to human dietary patterns than the human reference (Fig. S4c). These observations are consistent with dietary amino acid supply contributing to the trading strategy of human AArchetype A.

We further evaluated the trading strategy in zebrafish, fruit flies and mice by comparing pathway amino acid compositions with those of species-specific experimental diets^24-26^. In zebrafish and mice, Cluster 1, which contained central carbon metabolism-related pathways, exhibited the smallest median distances to the corresponding diet among the species-specific clusters (Fig. 4f). The same pattern was apparent across the additional zebrafish and mouse dietary formulations (Fig. S4d,e). In fruit flies, Cluster 3 was slightly closer to the standard diet than Cluster 1, the cluster most enriched for central carbon metabolism pathways. However, Cluster 3 also contained central carbon metabolism pathways, and both clusters were substantially closer to the diet than Cluster 2 (Fig. 4f). These results extend the association between central carbon metabolism and dietary amino acid matching beyond humans, providing further support for the proposed trading strategy across other species.

Together, these results support a “trading” strategy underlying AArchetype A. Central carbon metabolism enzymes, which originated in ancient metabolic networks and are evolutionarily conserved and highly expressed, may reduce their effective amino acid costs by relying on dietary amino acid supplies that already approximate their compositional requirements.

### Costly AArchetypes are associated with structural disorder

We next asked whether the high costs of AArchetypes B and D reflect additional biophysical or cellular constraints that require incorporation of high-cost amino acids. For instance, transmembrane domains of membrane proteins require hydrophobic amino acids, therefore shaping their amino acid composition^38, 39^. Membrane association partly explains the emergence of AArchetypes: AArchetypes C and D were strongly enriched for membrane proteins compared to AArchetypes A and B (Fig. 5a). However, the two high-cost AArchetypes involve both soluble (B) and membrane-associated (D) enzymes (Fig. 5a), indicating that costly enzymes exist within both localization contexts.

**Fig. 5.**
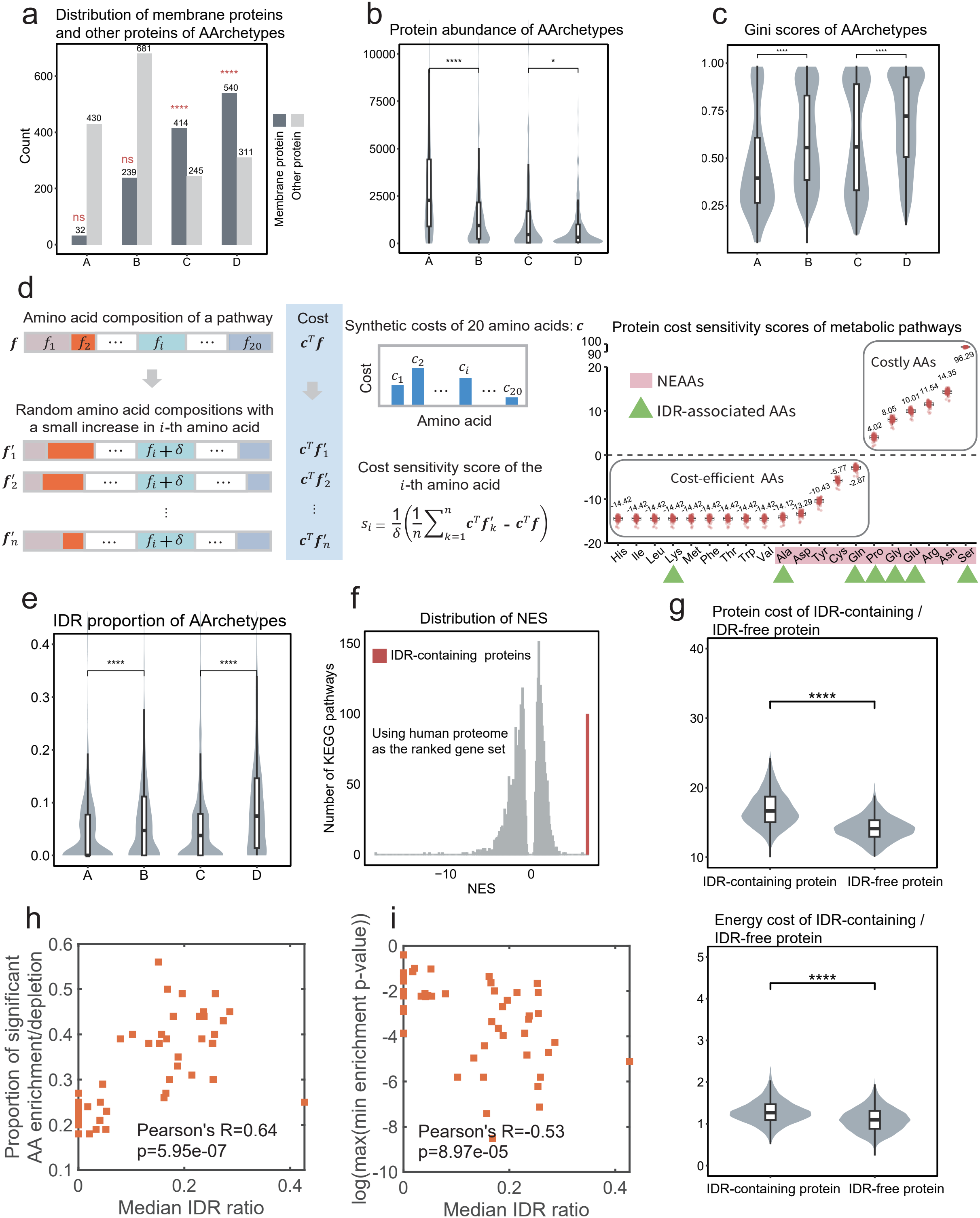
Costly AArchetypes are associated with structural disorder and phase separation. a: Counts of membrane proteins and other proteins in human AArchetypes. Statistical significance was evaluated using hypergeometric test. Significance: ns: p≥0.05; *: p<0.05; **: p<0.01; ***: p<0.001; ****: p<0.0001. b: Violin plot comparing protein abundance across the four human AArchetypes. Statistical significance was evaluated using Wilcoxon’s rank-sum test. Significance: ns: p≥0.05; *: p<0.05; **: p<0.01; ***: p<0.001; ****: p<0.0001. c: Distribution of Gini coefficients quantifying tissue specificity of protein abundance in AArchetypes. Statistical significance was evaluated using Wilcoxon’s rank-sum test. Significance: ns: p≥0.05; *: p<0.05; **: p<0.01; ***: p<0.001; ****: p<0.0001. d: Schematic overview of the calculation of protein cost sensitivity scores for human metabolic pathways (top) and the resulting scores (bottom). The numbers above each box plot represent the median value. e: Distribution of IDR proportion for enzymes in the human AArchetypes. Statistical significance was evaluated using Wilcoxon’s rank-sum test. Significance: ns: p≥0.05; *: p<0.05; **: p<0.01; ***: p<0.001; ****: p<0.0001. f: Counts of LLPS and Non-LLPS proteins in AArchetypes. Statistical significance was evaluated using hypergeometric test. g: Distribution of normalized enrichment scores (NES) from GSEA for gene sets enriched in costly human proteins. Gene sets include KEGG pathways, and curated sets of IDR- and LLPS-associated proteins. h: Distribution of protein cost (left) and energy cost (right) of LLPS/non-LLPS protein across the whole human proteome. Statistical significance was evaluated using Wilcoxon’s rank-sum test. Significance: ns: p≥0.05; *: p<0.05; **: p<0.01; ***: p<0.001; ****: p<0.0001.

Using quantitative proteomics data across human tissues (Methods), we found that enzymes within the two high-cost AArchetypes are expressed at lower abundance relative to their counterparts within the same localization group: B is lower than A among non-membrane enzymes, and D is lower than C among membrane-enriched enzymes (Fig. 5b). Moreover, protein abundances of enzymes in the costly AArchetypes B and D are tissue specific as reflected by higher Gini coefficients (Fig. 5c, Methods).

Together, these results suggest that enzymes with costly amino acid composition tend to reduce their realized cost by lowering overall abundance and restricting expression to specific tissues. However, such association alone does not explain the existence of costly AArchetypes, as adopting a less costly amino acid composition can always reduce the cost of enzymes regardless of the protein abundance. To identify features associated with these costly AArchetypes, we next sought to determine single amino acids contributing to increased cost of enzymes. By computing protein and energy cost sensitivity scores quantifying the change in pathway-level costs with a marginal compositional increase of each single amino acid (Fig. 5d, S5a, Methods), we identified a subset of amino acids associated with the costly AArchetypes, including proline, glycine, glutamate, arginine, asparagine, and serine (Fig. 5d). This set overlaps substantially (4 out of 5) with non-essential amino acids that are frequently enriched in intrinsically disordered regions (IDRs), including arginine, serine, glutamine, proline, and glutamate^40^ (Fig. 5d), establishing a connection between high synthetic cost and intrinsic structural disorder.

The conformational flexibility of IDRs supports diverse and often transient interactions with proteins and nucleic acids, which are frequently involved in a variety of dynamic biological processes such as signal transduction, transcriptional control^41^, and liquid-liquid phase separation (LLPS)^42^. For metabolic enzymes, involvement of IDRs can carry functional benefits by facilitating adaptation to different substrates, enhancing controllability of reaction kinetics, supporting moonlighting functions beyond catalysis^43^, and enabling dynamic, reversible compartmental organization^44-47^. Nevertheless, excessive or aberrant accumulation of IDR-containing proteins may disrupt cellular interaction networks through inappropriate molecular interactions, aberrant condensate formation, and pathological aggregation, with implications for neurodegeneration and cancer^48-50^. Consequently, abundances of many IDR-containing proteins are tightly controlled at low levels through mechanisms including transcription, translation, and protein degradation^51^. We therefore hypothesize that elevated biosynthetic costs of IDR-associated amino acids may impose a resource constraint on the accumulation of IDR-rich proteins, increasing the cellular cost of producing them at high abundance. Introduction of such resource constraint might mitigate the potentially harmful effects of excessive IDRs on cellular interaction networks. We found that for the membrane-unrelated AArchetypes A and B, enzymes in the higher-cost AArchetype B exhibit a higher IDR proportion (Fig. 5e) compared to those in the lower-cost AArchetype A. A similar association of higher cost with increased IDR content was observed when comparing the membrane-associated AArchetypes C and D (Fig. 5e), indicating a connection between structural disorder and high cost independent of enzyme localization in humans.

To test whether this coupling extends beyond metabolic enzymes, we performed gene set enrichment analysis (GSEA) to identify functional categories associated with high cost throughout all human proteins. Compared to all KEGG pathways, the set of IDR-containing proteins exhibited the highest normalized enrichment score (NES = 6.6), markedly exceeding those of all KEGG pathways, whose NES values were predominantly clustered around zero with a negative tail (Fig. 5f, S5b-e), indicating that involvement of IDR is the most pronounced feature of high-cost proteins. Direct proteome-wide comparisons further confirmed that IDR-containing proteins have significantly higher protein and energy costs than non-IDR-containing proteins (Fig. 5g). These results indicate that the association between high synthetic cost and enrichment of IDR is not limited to metabolic enzymes, but exists across all human proteins.

Furthermore, to determine whether the association between structural disorder and AArchetype emergence extends beyond humans, we compared IDR proportions among AArchetypes in all species classified as exhibiting AArchetype emergence (Fig. 2c). In 16 of the 17 species examined, at least one pair of AArchetypes differed significantly in IDR proportion, with *Amphimedon queenslandica* as the sole exception (Fig. S5). This widespread cross-species pattern suggests that structural disorder may be a recurrent contributor to AArchetype diversification across species. The median IDR ratio also showed significant correlations with the two metrics for AArchetype emergence: the proportion of significant AA enrichment/depletion (Pearson’s *R* = 0.64; Fig. 5h), and the log-transformed max–min enrichment *P* value (Pearson’s *R* = −0.53; Fig. 5i). The correlation for max-min enrichment p-value remained significant after adjusting for phylogeny, therefore linking structural disorder to the functional signatures of AArchetypes independent of shared phylogenetic history.

Taken together, the association between high synthetic cost and IDRs suggests an economic interpretation for why costly AArchetypes B and D persist. IDRs can be functionally valuable for enzymes with this capability but potentially hazardous at the entire cell level, therefore creating a negative externality. A plausible way for the cell to internalize this externality is to effectively raise the costs of IDR-associated amino acids, thereby making IDR-containing proteins intrinsically expensive to produce and favor regulatory strategies that limit their total burden. In this framework, AArchetypes B/D complement cost saving (C) and trading (A) as a third strategy: internalization of externality, where amino acids associated with negative externality are subject to enhanced synthetic costs.

### A mathematical model for IDR externality internalization by increasing synthetic cost

The coupling between high synthetic cost and IDR in the human AArchetypes motivates an economic interpretation: IDR can provide protein-level functional benefits, but excessive or dysregulated accumulation of IDR-containing proteins could impose a burden at the cellular level. Such interpretation is consistent with the observation that overexpression of IDR-rich proteins is associated with greater fitness costs in several CRISPR activation screening studies across species (Figure S7a, b). In economic theory, such a mismatch between local benefit and global burden is described by negative externality, which can be internalized through price-based instruments such as Pigouvian taxation. We therefore constructed a simplified mathematical model to examine whether elevated synthetic costs of IDR-related amino acids can act as an effective pricing constraint that restrains IDR level while preserving functional benefits (Fig. 6a, Methods). We used the model to examine how amino acid prices shape the optimal amino acid composition and protein abundance. We also performed a counterfactual simulation in which human proteins optimize their amino acid compositions under a pricing system that does not elevate the costs of IDR-related amino acids.

**Fig. 6.**
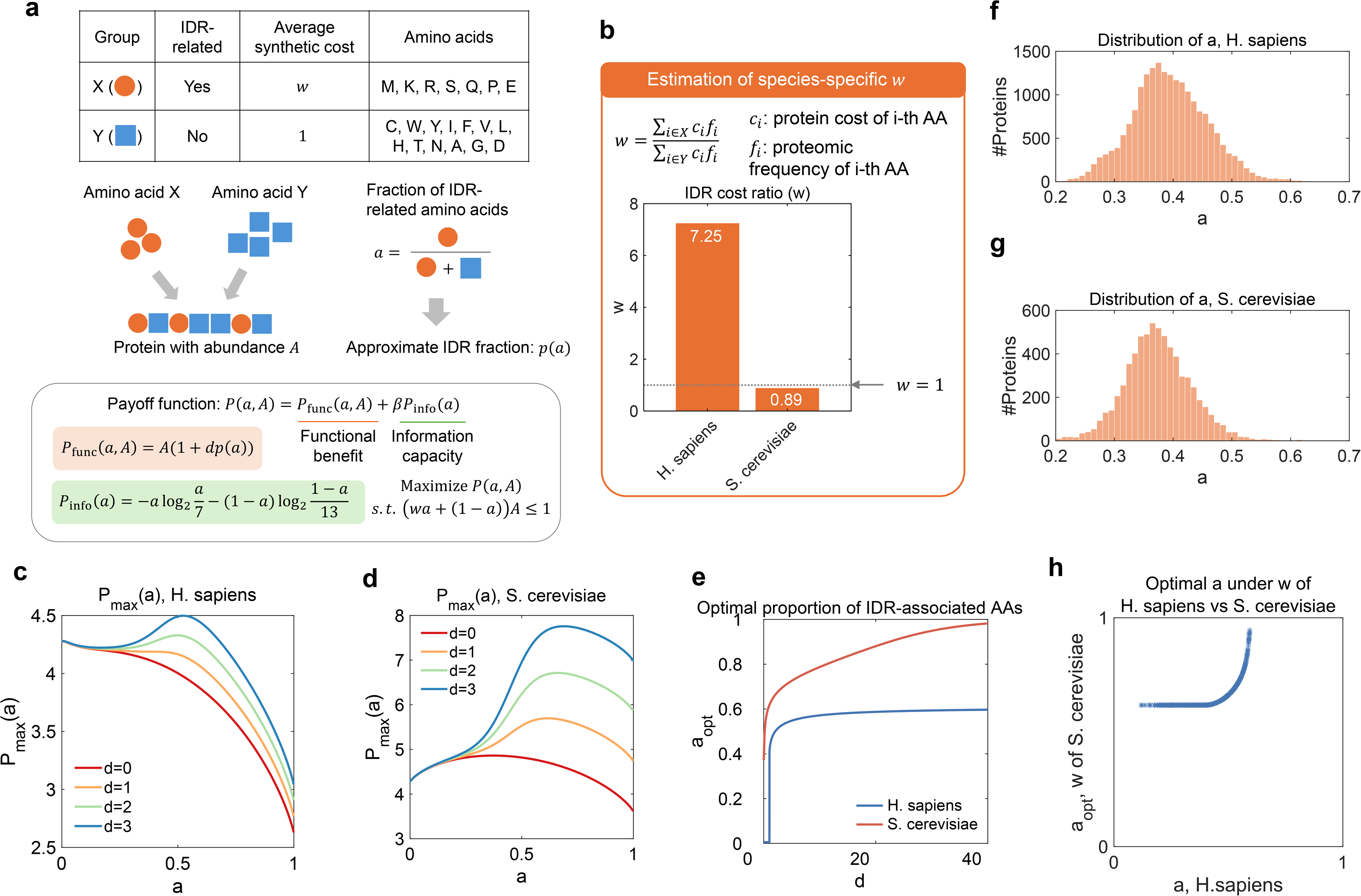
A mathematical model for IDR externality internalization by increasing synthetic cost. a: Schematic overview of the mathematical model. b: Algorithm for estimation of the species-specific IDR cost ratio *w* and bar plot showing the values of *w* in human (*H. sapiens*) and yeast (*S. cerevisiae*). c: Maximal payoff function *P*_max_*(a)* under varying *a* for human (*H. sapiens*). d: Maximal payoff function *P*_max_*(a)* under varying *a* for yeast (*S. cerevisiae*). e: Optimal solution of the fraction of IDR-related amino acids *a* under varying IDR payoff gain *d* under the human (*w = 7*.*25*) and yeast (*w = 0*.*89*) pricing system. f: Distribution of the fraction of IDR-related amino acids *a* in human proteins. g: Same as in (f) but for yeast proteins. h: Scatter plot comparing the actual fraction of IDR-related amino acids *a* and the simulated optimal *a* under the yeast-like amino acid pricing regime.

The model involves two groups of amino acids: an IDR-related amino acid group *x* and an IDR-unrelated amino acid group *Y*, that together forms a protein with abundance *A*. The protein determines its fraction of IDR-related amino acids, *a*, by maximizing a payoff function under the constraint of limited budget. Functional benefit of IDR increases with the fraction of IDR in the protein sequence *p(a)*. The relationship between fraction of IDR and fraction of IDR-related amino acids, *p(a)*, was obtained by fitting a logistic function to real data of human proteins (Methods, Figure S7c). The payoff function involves a functional benefit term related to the protein abundance A and an IDR-dependent functional gain proportional to *p(a)* (Fig. S7a, Methods), and an information capacity term quantifying the sequence complexity:

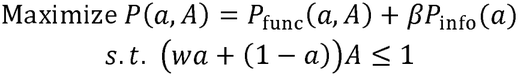

The model contains two IDR-related parameters: the IDR payoff gain *d* and the IDR cost ratio *w*. The IDR payoff gain *d* is a protein-specific parameter quantifying the increase in the payoff function contributed by one additional unit of IDR fraction. The IDR cost ratio *w* is a species-specific parameter estimated by comparing synthetic costs of IDR-related and IDR-unrelated amino acids (Fig. 6b, Methods). A value of *w* over 1 indicates a higher cost of IDR-related amino acids. IDR cost ratio *w* differs by one order of magnitude between human (*H. sapiens*) and yeast (*S. cerevisiae*): *w*=7.25 for human and 0.89 for yeast (Fig. 6b), indicating higher costs of IDR-related amino acids in human but not in yeast. As yeast proteins contain much lower fraction of IDR (median IDR fraction = 0.19) compared to human proteins (median IDR fraction = 0.51), we assumed that a yeast protein with median *a* (*a = 0*.*37*) has zero IDR payoff gain (*d= 0*) and estimated the parameter *β*accordingly (Fig. S7d, e, Methods).

The difference in IDR cost ratio *w* between human and yeast results in distinct modes of amino acid compositional optimization. To illustrate such distinction, we computed the maximal payoff function under varying *a, P*_max_*(a)*, for both human (*w*=7.25, Fig. 6c) and yeast (*w*=0.89, Fig. 6d). For human, the relationship between *P*_max_*(a)* and *a* depends largely on the IDR payoff gain parameter *d*: *P*_max_*(a)* reaches its maximum at *a= 0* for smaller values of d, and peaks at *a ≈ 0*.*5* for larger *d* (Fig. 6c,e). For yeast, the value of *a* that maximizes *P*_max_*(a)* approaches 1 at larger *d*. This is because in the absence of the penalty by costly IDR-related amino acids, the only factor that prevents *a* from becoming 1 is the low information capacity for *a* near 1 (Fig. S7e), which can be compensated by the protein abundance *A* (Fig. S7f) and IDR reward *dp(a)* (Fig. S7a) that both increase with *a*. Such difference suggests that the higher costs of IDR-related amino acids in the human model create a trade-off between increased IDR reward and decreased protein abundance when *a* increases, allowing regulation of *a*_opt_ even under strong functional benefit of IDR.

To connect the model predictions to real data of human proteins, we examined the distributions of *a* in human (Fig. 6f) and yeast (Fig. 6g) proteins. The range of *a* in real human proteins aligns well with the model-predicted range of *a*_opt_, further supporting the model and allowing direct inference of the IDR payoff gain *d* for almost all human proteins (20291 out of 20402, Fig. 6h, Methods). We therefore inferred IDR payoff gain *d* for those human proteins, and performed a counterfactual simulation in which human proteins retain their inferred IDR payoff gains but are optimized under the yeast-like pricing regime. In this scenario, optimal *a* for all human proteins (minimal value across all proteins = 0.62) becomes greater than the 99% quantile of real *a* in human proteins (99% quantile = 0.57, Fig. 6h). This outcome increases protein-level payoff but would dramatically expand the burden of IDR-related potentially harmful effects at the cellular level, consistent with a scenario of negative externality. Thus, the model provides a quantitative illustration of how the elevated synthetic costs of IDR-related amino acids can internalize IDR-related negative externalities.

## Discussion

In this study, we reveal economic strategies underlying the diversification of amino acid composition of metabolic enzymes into AArchetypes, i.e. functionally related clusters of metabolic enzymes that share similar amino acid compositions. Catalyzing specific biochemical reactions, the primary goal of metabolic enzymes, does not require the amino acid composition to be strictly fixed^52^. Therefore, like how Carl Jung’s archetypes reflect conserved, universal patterns in the psychological architecture of all human beings, the emergence of AArchetypes in metabolic enzymes indicates previously unexplored, catalysis-independent forces that shape amino acid sequences of enzymes, such as evolutionary history, environmental conditions, and economic strategies in complex multicellular organisms.

Across a taxonomically broad set of species, AArchetype signals tended to strengthen with the number of protein families, a proxy for organismal complexity. The observation suggests, unlike the principle of synthetic cost minimization in unicellular organisms such as bacteria and yeast, minimization of synthetic cost no longer governs the global amino acid composition of the proteome of complex multicellular species. Using human metabolic enzymes as a model system for proteome in multicellular life, our analyses define two alternative economic strategies: trading, in which dietary amino acid composition matches that of conserved and abundant central carbon metabolic enzymes; and internalization of negative externality, in which IDR-associated amino acids exhibit enhanced synthetic costs to prevent excessive accumulation of disordered proteins.

The emergence of these two economic strategies reflects fundamental differences between unicellular and multicellular organisms. Unicellular organisms and multicellular organisms differ intrinsically in strategy of nutrient acquisition. While unicellular organisms often rely on *de novo* biosynthesis from fluctuating, opportunistic supply of small molecule nutrients to meet their metabolic demands, nutrient supply in multicellular organisms becomes increasing buffered and internalized through organism-level systems for digestion, absorption, and long-range transport between tissues. Such systems exist across the kingdom of multicellular life, spanning digestion and circulation systems in animals, root, xylem, and phloem in plants, and interconnected mycelial networks in fungi^53^. This transition transforms amino acids from stochastic environmental inputs into regulated resources, enabling a shift from purely biosynthetic cost minimization toward a trading-based strategy in which external supply can partially substitute for endogenous production.

Second, the enhanced synthetic costs of IDR-related amino acids in human metabolism suggests a strategy of IDR externality internalization. IDRs are recognized as functional structural elements capable of mediating flexible linkers, multivalent and reversible interactions, and dense post-translational regulation^54-56^; nevertheless, dysfunctional IDRs are associated with a variety of pathologies. Negative externality, a concept borrowed from economics, describes the harmful impacts that actions of business firms have on others. An example is a factory that pollutes the air while making its products, as the factory benefits from selling the products while society suffers from the pollution. In economic theory, negative externalities are commonly internalized through three strategies: price-based instruments, quantity control, and regulatory mechanisms. Price-based instruments, such as Pigovian tax, increase the private cost associated with activities generating social harm^57^. Quantity-based instruments restrict total allowed scale of such activities, while regulatory mechanisms provide avenues for elimination of the harmful effects, limiting long-term accumulation and systemic risk. Both quantity-based and regulatory strategies have clear counterparts in the cellular control of intrinsic disordered proteins. Dosage-sensitive genes tend to encode IDR-containing proteins^58-60^, suggesting that the total allowed amount of disordered proteins is tightly controlled by selective pressure disfavoring their amplification; IDR-containing proteins have shorter half-lives^61^, limiting their steady state concentration. These observations imply the necessity of controlling excessive intrinsic disordered proteins and point to abundance limitation as a quantity-control mechanism for managing IDR-associated risks. Together with the Pigovian taxation of IDR-related amino acids proposed in our theory, these mechanisms form a multi-layer economic system controlling the negative externality associated with IDR.

Although experimental determination of synthetic costs is challenging, our theory generates predictions that can be experimentally tested in future studies. For instance, disruption of the Pigovian taxation of IDR-related amino acids in human cells, such as expression of high-activity mutants of enzymes in biosynthesis of costly amino acids, could lead to observable changes in the fitness and phenotype of cells. Changes in the composition of dietary amino acid intake could also perturb the “trading” system, therefore reprogramming central carbon metabolism catalyzed by enzymes predicted to rely more on the trading strategy. It is worth noting that our previous large-scale analysis of amino acid composition of human foods^28^ shows that the relative ordering of amino acid abundance in human foods is remarkably robust, supporting the idea that dietary intake of amino acids is constrained to match the composition of conserved central carbon metabolic enzymes.

In summary, our study reveals previously unexplored complexity in the amino-acid economics of multicellular life. In these organisms, the emergence of long-range transport, dietary buffering, and IDR-dependent interactions gives rise to a richer economic landscape in which production, trading, and regulation coexist. Hence, the emergence of AArchetypes highlights a transition from a cost-driven market to a governed economy as a fundamental hallmark of complex life.

## Data availability

All datasets used and generated in this study are available at the GitHub page: https://github.com/Alec-Yeung/AArchetypes.

## Code availability

All code and scripts generated in this study are available at the GitHub page: https://github.com/Alec-Yeung/AArchetypes.

## Author contributions

K.Y. and Z.D. designed the study and wrote the manuscript. K.Y. and J.W. performed the AArchetype-related analyses. K.S. retrieved and processed the human and yeast proteomics datasets. H.Z. and Y.L. analyzed amino acid sequences of human metabolic enzymes. Z.D. developed the mathematical model for IDR externality internalization. All authors have read and approved the final manuscript.

## Acknowledgements

This research was supported by the National Natural Science Foundation of China (12371489 to ZD and 12205012 to YL), National Key Research and Development Program of China (2021YFA0911300 to ZD), Shenzhen Medical Academy of Research and Translation (D2401022 to ZD), and Basic and Applied Basic Research Foundation of Guangdong Province (2025A1515012923 to YL). The authors thank Prof. Cong Yu, Prof. Kaige Yan, and Prof. Xiandeng Wu for helpful discussions.

## Notes

### Competing Interest Statement

The authors have declared no competing interest.

### Summary of Updates

Revised cross-species analysis of AArchetypes, IDR-related analysis, and added dietary matching analysis for more species.

