## Supplementary Figures for "Cost-saving, trading, and internalization of externality: three economic strategies underlying amino acid archetypes of human metabolic enzymes"

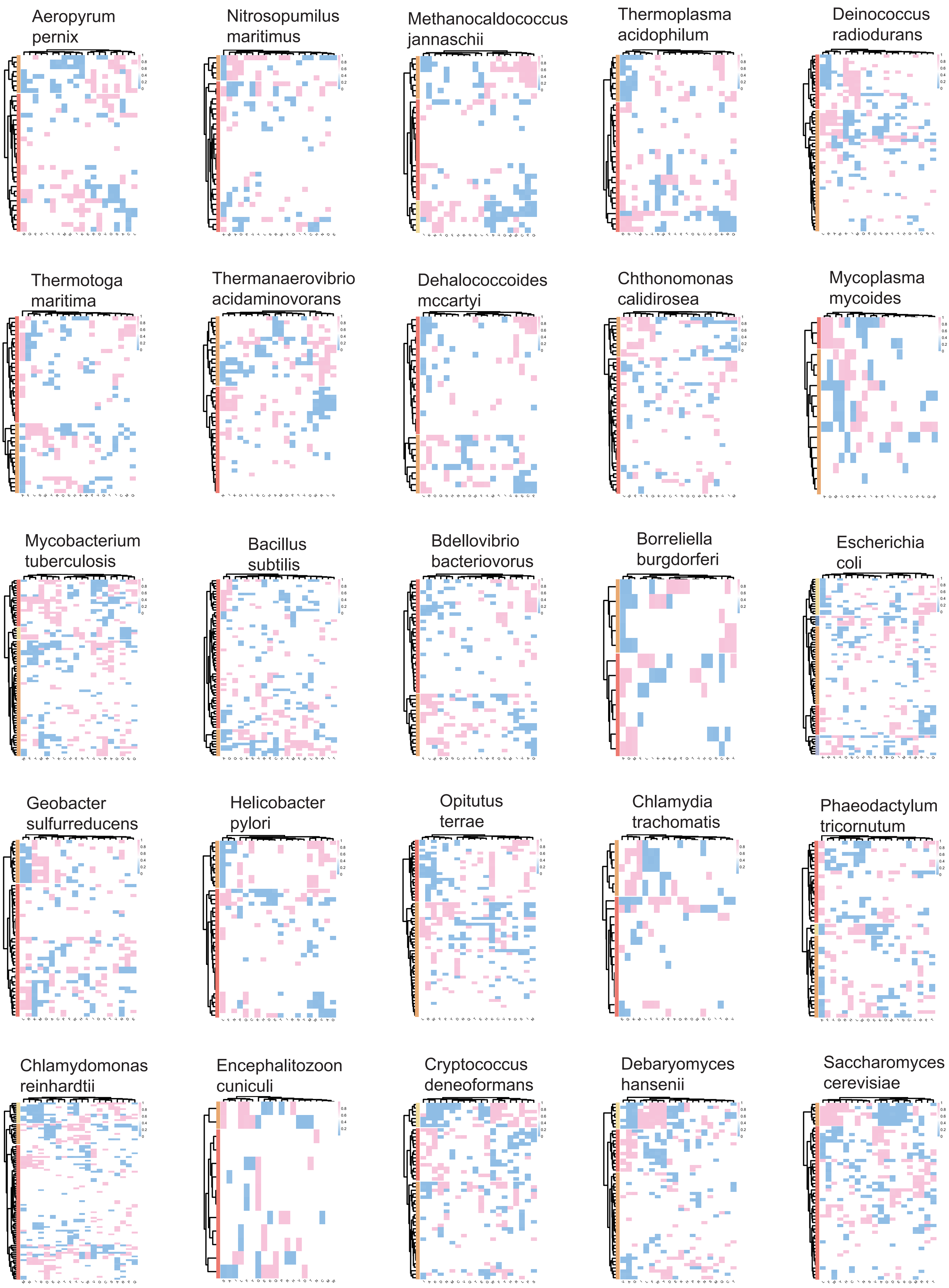

Aeropyrum pernix
Nitrosopumilus maritimus
Methanocaldococcus jannaschii
Thermoplasma acidophilum
Deinococcus radiodurans
Thermotoga maritima
Thermanaerovibrio acidaminovorans
Dehalococcoides mccartyi
Chthonomonas calidirosea
Mycoplasma mycoides
Mycobacterium tuberculosis
Bacillus subtilis
Bdellovibrio bacteriovorus
Borreliella burgdorferi
Escherichia coli
Geobacter sulfurreducens
Helicobacter pylori
Opitutus terrae
Chlamydia trachomatis
Phaeodactylum tricornutum
Chlamydomonas reinhardtii
Encephalitozoon cuniculi
Cryptococcus deneoformans
Debaryomyces hansenii
Saccharomyces cerevisiae

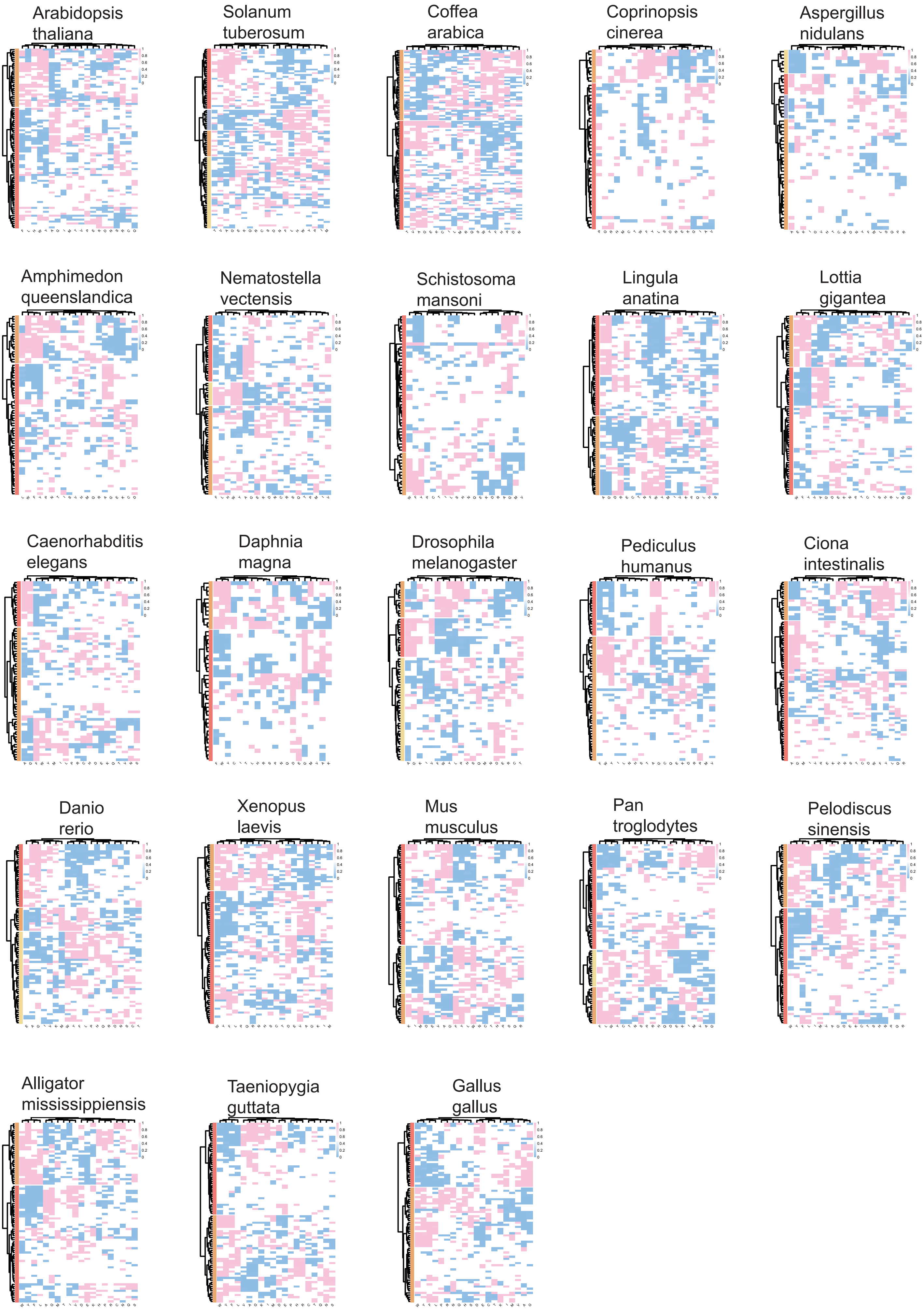
Arabidopsis thaliana
Solanum tuberosum
Coffea arabica
Coprinopsis cinerea
Aspergillus nidulans
Amphimedon queenslandica
Nematostella vectensis
Schistosoma mansoni
Lingula anatina
Lottia gigantea
Caenorhabditis elegans
Daphnia magna
Drosophila melanogaster
Pediculus humanus
Ciona intestinalis
Danio rerio
Xenopus laevis
Mus musculus
Pan troglodytes
Pelodiscus sinensis
Alligator mississippiensis
Taeniopygia guttata
Gallus gallus

a

Protein cost and energy cost of AAs

| AA | AA abbreviation | Protein cost | Energy cost |
|---|---|---|---|
| Ala | A | 0.3 | 0 |
| Cys | C | 8.7 | 0 |
| Asp | D | 1.1 | 0 |
| Glu | E | 24.4 | 0 |
| Phe | F | 0 | 0 |
| Gly | G | 22.5 | 0 |
| His | H | 0 | 0 |
| Ile | I | 0 | 0 |
| Lys | K | 0 | 0 |
| Leu | L | 0 | 0 |
| Met | M | 0 | 0 |
| Asn | N | 28.8 | 5 |
| Pro | P | 18.5 | 0 |
| Gln | Q | 11.6 | 5 |
| Arg | R | 26 | 15 |
| Ser | S | 110.8 | 0 |
| Thr | T | 0 | 0 |
| Val | V | 0 | 0 |
| Trp | W | 0 | 0 |
| Tyr | Y | 4 | 0 |

b

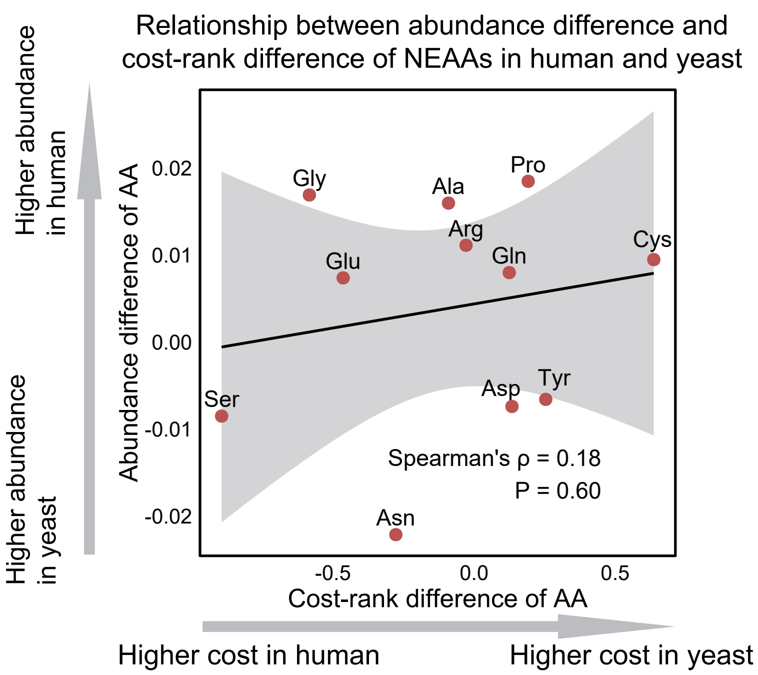

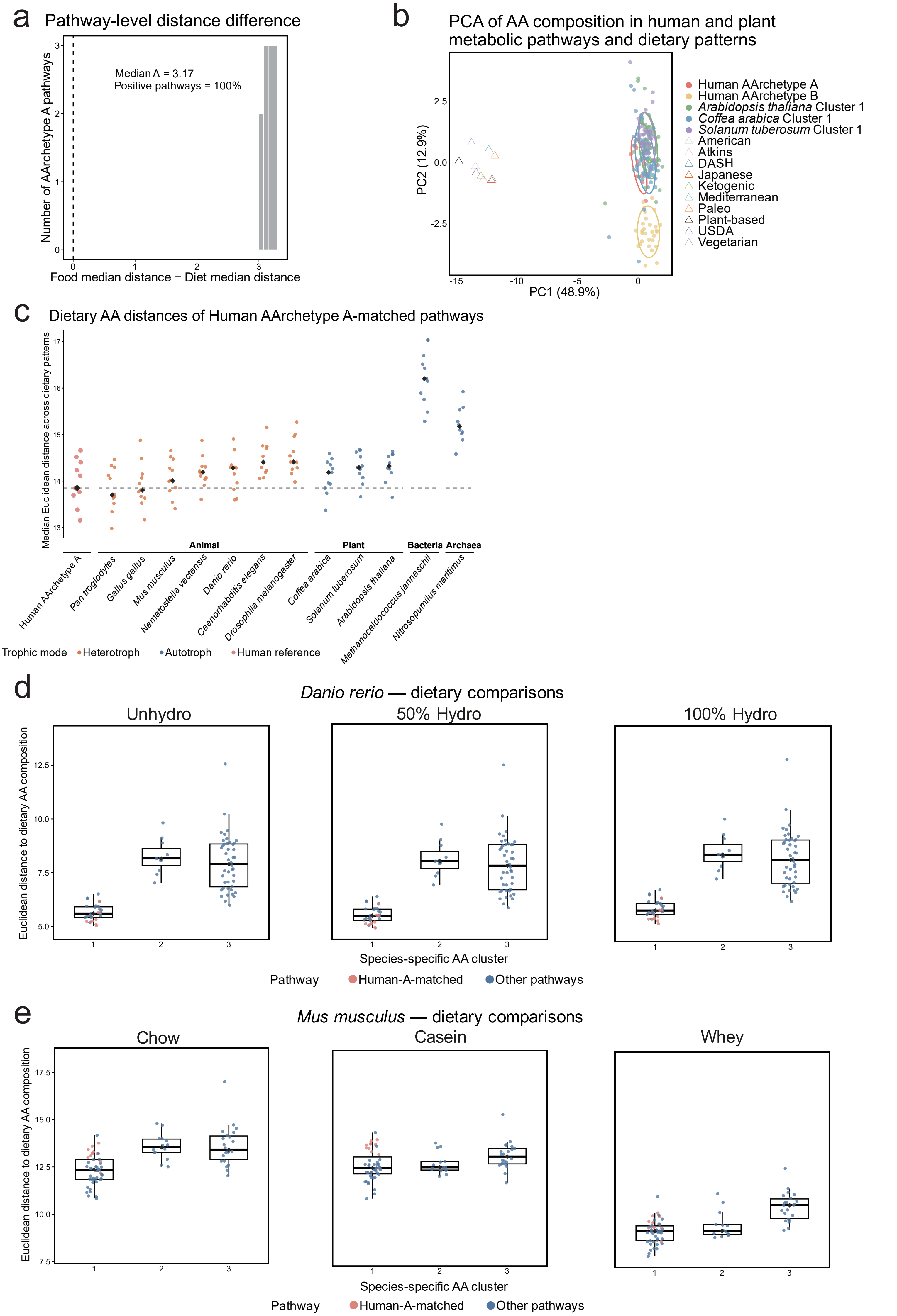
a
Pathway-level distance difference
Median Δ = 3.17
Positive pathways = 100%
Number of AArchetype A pathways
Food median distance − Diet median distance
b
PCA of AA composition in human and plant metabolic pathways and dietary patterns
PC1 (48.9%)
PC2 (12.9%)
Human AArchetype A
Human AArchetype B
Arabidopsis thaliana Cluster 1
Coffea arabica Cluster 1
Solanum tuberosum Cluster 1
American
Atkins
DASH
Japanese
Ketogenic
Mediterranean
Paleo
Plant-based
USDA
Vegetarian
c
Dietary AA distances of Human AArchetype A-matched pathways
Median Euclidean distance across dietary patterns
Animal
Plant
Bacteria
Archaea
Human AArchetype A
Pan troglodytes
Gallus gallus
Mus musculus
Nematostella vectensis
Danio rerio
Caenorhabditis elegans
Drosophila melanogaster
Coffea arabica
Solanum tuberosum
Arabidopsis thaliana
Methanocaldococcus jannaschii
Nitrosopumilus maritimus
Trophic mode
Heterotroph
Autotroph
Human reference
d
Danio rerio — dietary comparisons
Unhydro
50% Hydro
100% Hydro
Euclidean distance to dietary AA composition
Species-specific AA cluster
Pathway
Human-A-matched
Other pathways
e
Mus musculus — dietary comparisons
Chow
Casein
Whey
Euclidean distance to dietary AA composition
Species-specific AA cluster
Pathway
Human-A-matched
Other pathways

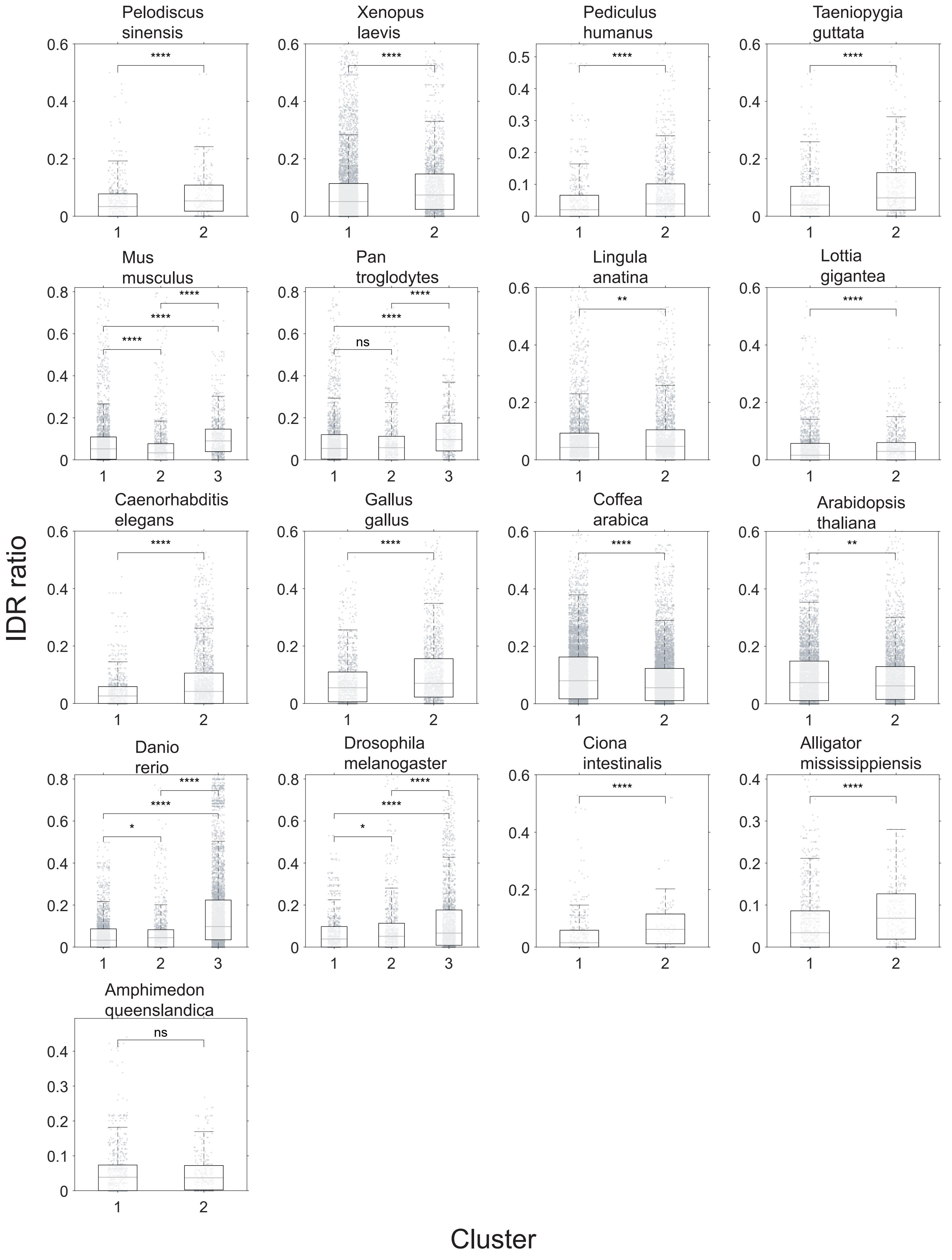
IDR ratio
Cluster
Pelodiscus sinensis
Xenopus laevis
Pediculus humanus
Taeniopygia guttata
Mus musculus
Pan troglodytes
Lingula anatina
Lottia gigantea
Caenorhabditis elegans
Gallus gallus
Coffea arabica
Arabidopsis thaliana
Danio rerio
Drosophila melanogaster
Ciona intestinalis
Alligator mississippiensis
Amphimedon queenslandica
ns

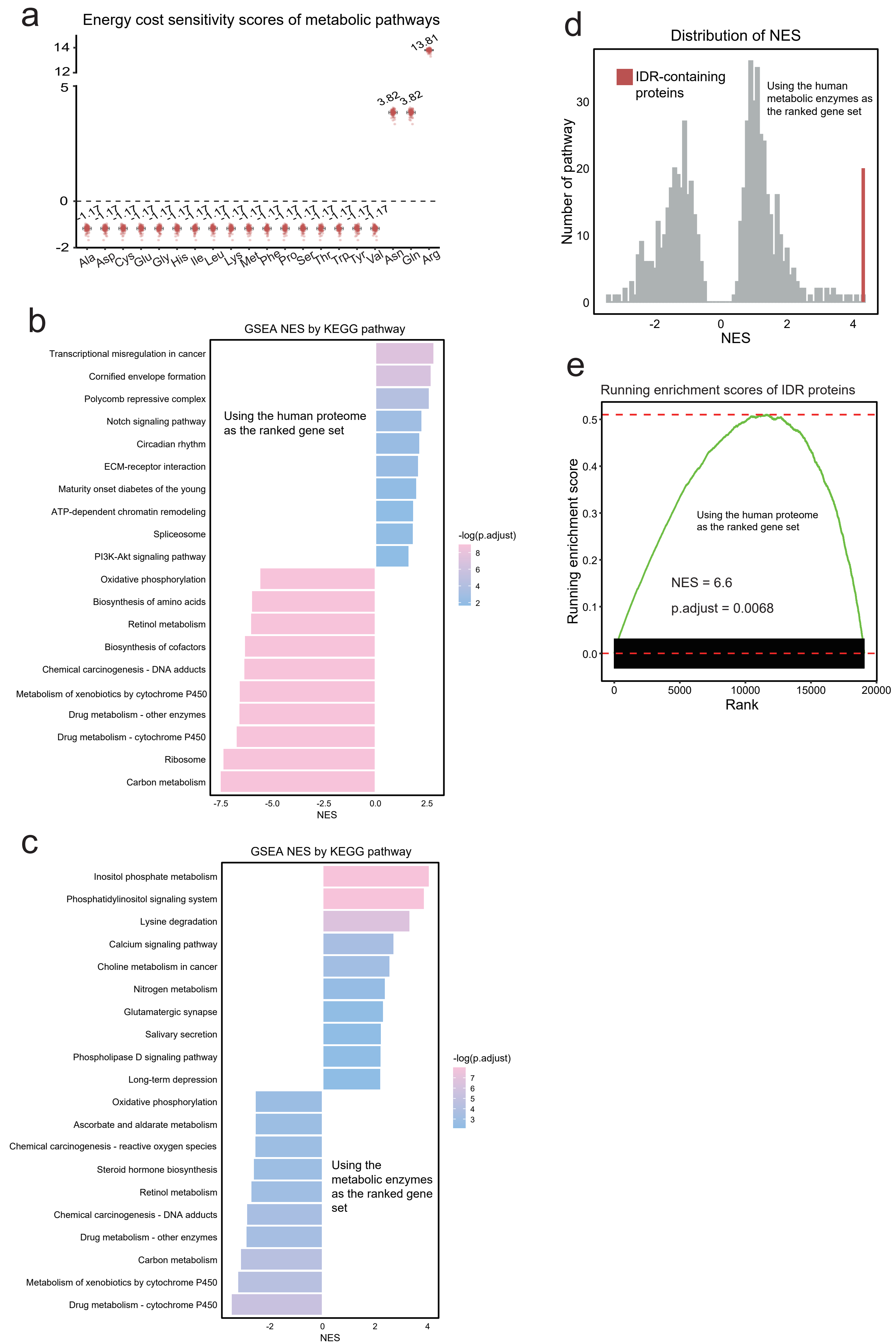

a
Energy cost sensitivity scores of metabolic pathways
13.81
3.82
3.82
-1.17
Ala Asp Cys Glu Gly His Ile Leu Lys Met Phe Pro Ser Thr Trp Tyr Val Asn Gln Arg
b
GSEA NES by KEGG pathway
Using the human proteome as the ranked gene set
Transcriptional misregulation in cancer
Cornified envelope formation
Polycomb repressive complex
Notch signaling pathway
Circadian rhythm
ECM-receptor interaction
Maturity onset diabetes of the young
ATP-dependent chromatin remodeling
Spliceosome
PI3K-Akt signaling pathway
Oxidative phosphorylation
Biosynthesis of amino acids
Retinol metabolism
Biosynthesis of cofactors
Chemical carcinogenesis - DNA adducts
Metabolism of xenobiotics by cytochrome P450
Drug metabolism - other enzymes
Drug metabolism - cytochrome P450
Ribosome
Carbon metabolism
NES
-log(p.adjust)
c
GSEA NES by KEGG pathway
Inositol phosphate metabolism
Phosphatidylinositol signaling system
Lysine degradation
Calcium signaling pathway
Choline metabolism in cancer
Nitrogen metabolism
Glutamatergic synapse
Salivary secretion
Phospholipase D signaling pathway
Long-term depression
Oxidative phosphorylation
Ascorbate and aldarate metabolism
Chemical carcinogenesis - reactive oxygen species
Steroid hormone biosynthesis
Retinol metabolism
Chemical carcinogenesis - DNA adducts
Drug metabolism - other enzymes
Carbon metabolism
Metabolism of xenobiotics by cytochrome P450
Drug metabolism - cytochrome P450
Using the metabolic enzymes as the ranked gene set
NES
-log(p.adjust)
d
Distribution of NES
IDR-containing proteins
Using the human metabolic enzymes as the ranked gene set
Number of pathway
NES
e
Running enrichment scores of IDR proteins
Running enrichment score
Using the human proteome as the ranked gene set
NES = 6.6
p.adjust = 0.0068
Rank

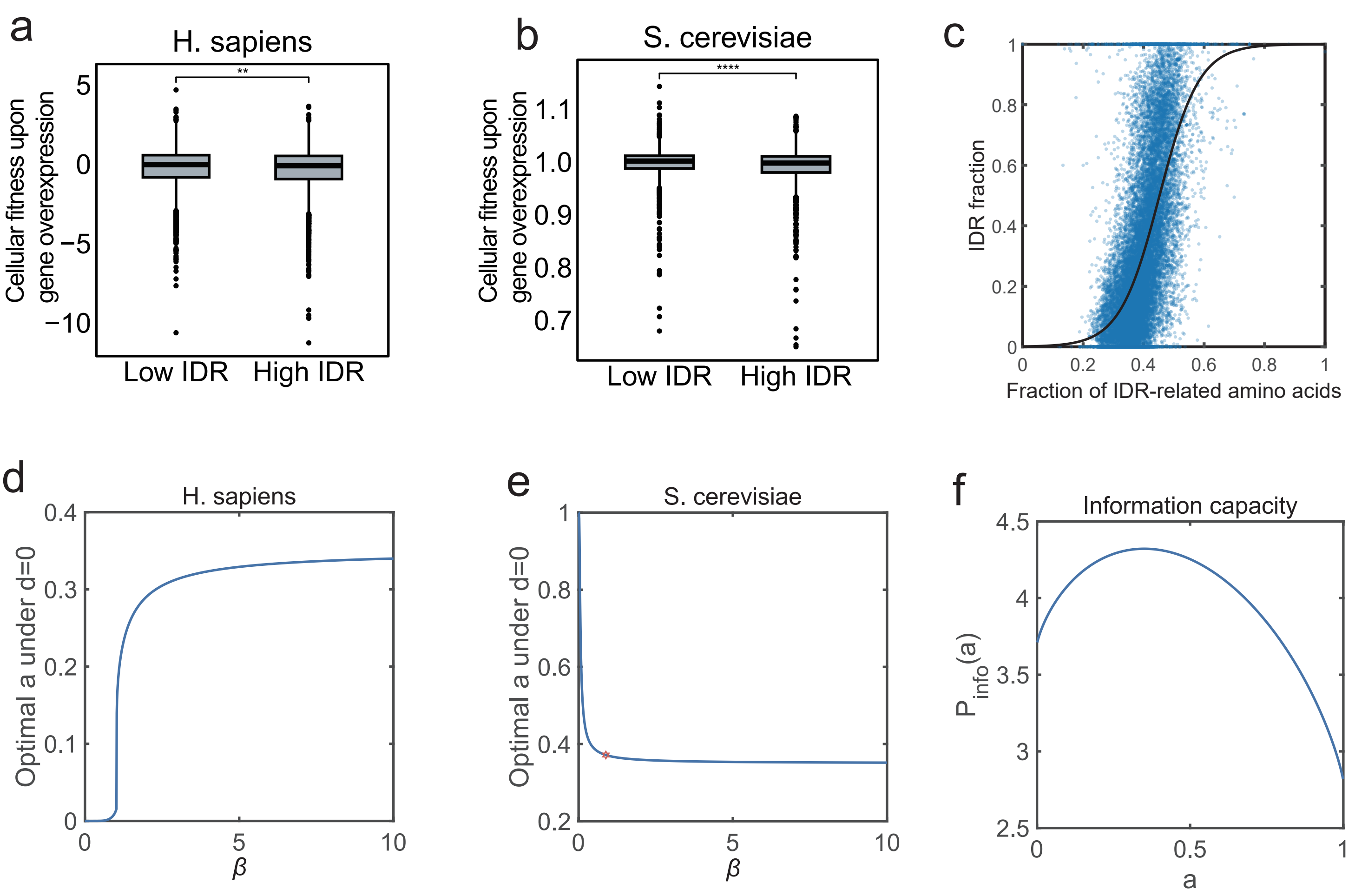

a
H. sapiens
Cellular fitness upon gene overexpression
**
Low IDR
High IDR
b
S. cerevisiae
Cellular fitness upon gene overexpression
****
Low IDR
High IDR
c
IDR fraction
Fraction of IDR-related amino acids
d
H. sapiens
Optimal a under d=0
β
e
S. cerevisiae
Optimal a under d=0
β
f
Information capacity
P_info(a)
a
